# Pluripotent stem cell derived models of neurological diseases reveal early transcriptional heterogeneity

**DOI:** 10.1101/2020.12.02.398263

**Authors:** Matan Sorek, Walaa Oweis, Malka Nissim-Rafinia, Moria Maman, Shahar Simon, Cynthia C. Hession, Xian Adiconis, Sean K. Simmons, Neville Sanjana, Xi Shi, Congyi Lu, Jen Q. Pan, Xiaohong Xu, Mahmoud A. Pouladi, Lisa M. Ellerby, Feng Zhang, Joshua Z. Levin, Eran Meshorer

## Abstract

**Background:** Many neurodegenerative diseases (NDs) develop only later in life, when cells in the nervous system lose their structure or function. In genetic forms of NDs, this late onset phenomenon remains largely unexplained.

**Results:** Analyzing single cell RNA sequencing (scRNA-seq) from Alzheimer’s disease (AD) patients, we find increased transcriptional heterogeneity in AD excitatory neurons. We hypothesized that transcriptional heterogeneity precedes ND pathologies. To test this idea experimentally, we used juvenile forms (72Q; 180Q) of Huntington’s disease (HD) iPSCs, differentiated them into committed neuronal progenitors, and obtained single cell expression profiles. We show a global increase in gene expression variability in HD. Autophagy genes become more stable, while energy and actin-related genes become more variable in the mutant cells. Knocking-down several differentially-variable genes resulted in increased aggregate formation, a pathology associated with HD. We further validated the increased transcriptional heterogeneity in CHD8^+/-^ cells, a model for autism spectrum disorder.

**Conclusions:** Overall, our results suggest that although NDs develop over time, transcriptional regulation imbalance is present already at very early developmental stages. Therefore, an intervention aimed at this early phenotype may be of high diagnostic value.

## Background

While, with some exceptions, all cells in the adult organism are genetically identical, they are not the same. This is true even in seemingly identical cells at the same developmental stage and at the same tissue. In addition to cyclic processes such as the cell cycle, cellular heterogeneity may arise as a result of intrinsic noise in the biochemical processes that operate within the cells. As the number of molecules that take part in many of these processes is relatively small, they become stochastic by nature [1].

One of the most extensively studied of those is the transcriptional machinery in the cell. The mRNA level of a specific gene in the cell typically depends on the product of only 1-2 DNA templates. Moreover, it is generally accepted that the transcription of most genes involves stochastic periods of transcriptional bursts (also called ‘on state’) during which mRNA is produced [2–4], rather than constitutively being transcribed. These bursty dynamics further increase the uncertainty in the number of mRNA molecules.

Interestingly, in the nervous system, damage to the cells may result in a change in the variability of mRNA levels. For example, during *C. elegans* development, the left Q neuroblast migrates posteriorly, a decision that is based on *Mab5* expression. A triple KO for three genes, which directly control *Mab5* expression, does not change the average expression of *Mab5* [5]. Instead, the distribution of the number of *Mab5* transcripts in the triple KO becomes more variable and results in more dispersed migratory distance, reflecting the impaired feedback control that is responsible for the robust expression. This raises the idea that transcriptional variability might contribute to the pathology of neurological disorders, such as Alzheimer’s disease (AD) or Huntington’s disease (HD).

HD is an autosomal dominant genetic neurodegenerative disorder (ND). it is caused by a repeat expansion in the *HTT* gene. The normal *HTT* gene contains, in its first exon, a coding sequence of a trinucleotide repeat of CAG/CAA (encoding the amino acid glutamine, denoted as Q), thus resulting in a protein that contains a polyglutamine (polyQ) tract. Both the normal and the mutant HTT proteins are expressed ubiquitously in all tissues. When the repeat length exceeds a threshold of 39 repeats, this results in the complete penetrance of the disease. Interestingly, the repeat length in healthy subjects, as well as in other primates, is much larger compared to the mouse orthologue, as is also the case in other polyQ-related diseases [6].

Given the ubiquitous and early expression of the mutant HTT protein [7], the reasons for the years-long delay in disease onset are not entirely clear. Under the general assumption that the symptoms of neural disorders are the result of broad enough neuronal malfunction, two extreme scenarios may explain this process. The first is the deterministic model. In this model, the WT and mutant cells both function normally at early development. However, they have different lifecycle trajectories; for example, the mutant cells, and not the WT cells, may gradually accumulate aggregates. Over time, the mutant cells become more and more diverse, until they are no longer functional, leading to cell death and disease onset. In contrast, in the stochastic model, both WT and mutant cells behave similarly. However, mutant cells cope with the cellular damage incurred as a result of the mutation, which comes at the expense of efficient self-regulation that maintains stable behavior. Therefore, over time, although both WT and mutant cells may stop functioning properly, the chances of a mutated cell reaching a ‘disease state’ are much higher compared to a WT cell. As a consequence, after sufficient time, a large enough fraction of mutated cells will stop functioning properly and initiate the symptoms of the disease.

Estimating the contribution of the stochastic hypothesis to the disease requires the quantification of the distribution of mRNA levels among cells. However, so far most of the studies on NDs have used bulk cell populations. While this allows picking up a global picture of the disease state, the details at the single cell level remain concealed. Recently however, several studies compared brains from AD patients to controls at the single cell level [8,9], providing us with an opportunity to test transcriptional heterogeneity in neurodegenerative conditions, and to search for evidence for the stochastic model in the transcriptional context.

We first analyzed available datasets of single cell (sc)RNA-Seq from AD brains [8,9], which indeed demonstrated evidence for increased transcriptional heterogeneity in adult neurons. However, our analysis also emphasized the need for controlled cellular systems, and for using Smart-seq2, which provides superior depth and consistency. In addition, the AD cells represent an advanced stage of the disease, and therefore do not provide information on transcriptional variability prior to disease onset. In order to test the idea of early transcriptional heterogeneity in NDs, we took advantage of pluripotent late-onset disease models, where we can measure the transcriptional signature of the ‘young’ cells, preceding any disease-related phenotypes. To this end, we used an induced pluripotent stem cell (iPSC) model of HD, making use of juvenile (72Q and 180Q) mutations, along with their corresponding isogenic controls [10,11]. We differentiated the iPSCs from these two isogenic systems towards neural progenitor cells (NPC), isolated single cells by Fluorescence-activated cell sorting (FACS), and applied scRNA-Seq using a modified Smart-seq2 protocol [12,13].

We show that the mutant HD cells are more heterogeneous compared to their corrected counterparts. This was also the case for isogenic CHD8^+/+^ and CHD8^+/-^ cells, a cellular model for autism spectrum disorder (ASD), consistent with our hypothesis that damaged neural cells have an increased cellular variability. Furthermore, by knocking down several differentially variable genes and analyzing polyQ aggregates in HEK293 and in iPSC-derived neuronal progenitor cells, we provide evidence that these changes are phenotypically relevant. Overall, we provide *in vivo* (human brain samples) and *in vitro* (human isogenic iPSCs) evidence for the stochastic model of gene expression in NDs.

## Results

To test whether transcriptional heterogeneity is observed in the adult, ND brain, we analyzed single-cell data obtained from the prefrontal cortex of 24 healthy and 24 patients with Alzheimer disease (AD) pathology. Although the data consist of a very large number (~80,000) of cells, the number of reads per cell was extremely variable ranging from ~200 to ~25,000 and the number of expressed genes between ~200 and 7,500. This wide range introduces biases that limit the ability to quantitatively compare the variability in gene expression levels between subjects. To account for the potential bias that can result from the difference between cells sequencing depths in the different conditions, we quantified the heterogeneity of each subject based only on the most highly expressed genes (see Methods). We validated that the contribution of the different neuronal subtypes was similar in AD and healthy subjects (p-value > 0.6, Kolmogorov-Smirnov test). Using this approach, we found that neurons, and specifically excitatory neurons, were significantly more heterogeneous in patients compared to healthy subjects, (Figure 1A-B and S1A-B), supporting our hypothesis. Other cell types, including astrocytes, oligodendrocyte progenitor cells and oligodendrocytes, also showed a similar trend albeit, mostly due to the small cell numbers, not significant (Figure S1C-F).

**Figure 1.**
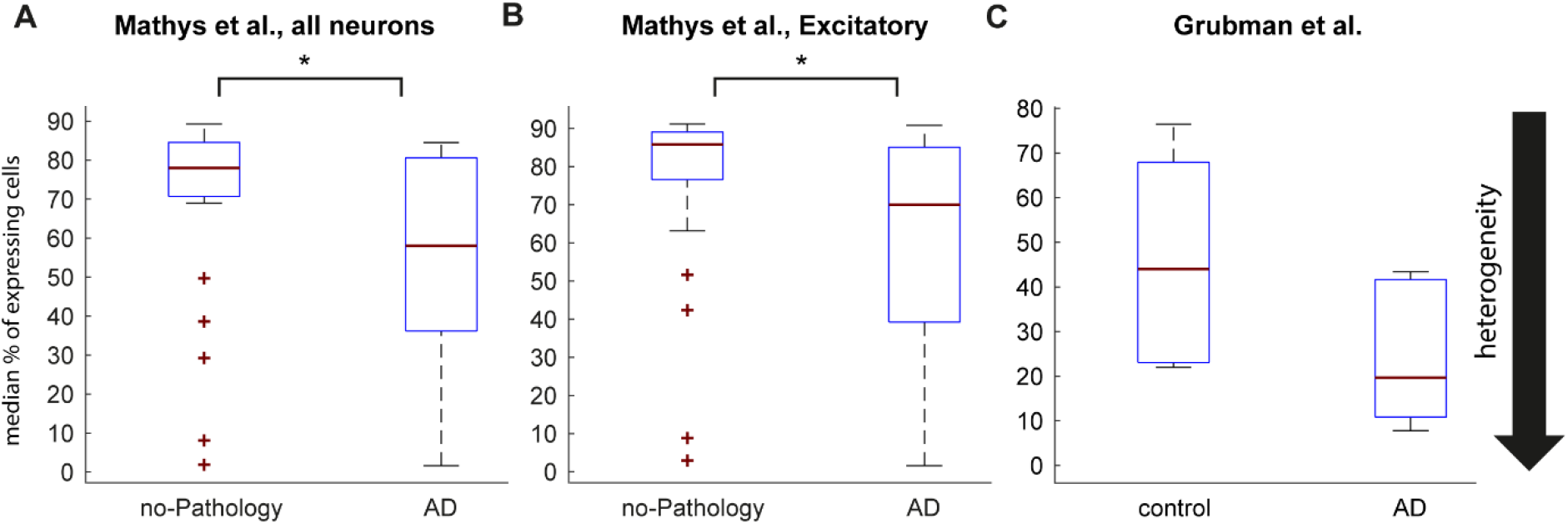
Adult neurons show increased transcriptional heterogeneity. (A-B) Boxplots showing the median percent of expressing cells of the highest 200 expressed genes in adult neurons from Mathys et al. (A, median percentages are 78% and 58% in healthy and AD subjects, respectively p = 0.0155, one-sided Wilcoxon rank-sum test) and in adult excitatory neurons (B, median percentages are 85.8% and 70% in healthy and AD subjects, respectively. p = 0.0073, one-sided Wilcoxon rank-sum test). n=24 healthy and n=24 AD subjects. FDR-corrected p-values remain significant with p=0.029 and 0.0312, respectively. (C) Same for the 100 highest expressed genes in neurons from Grubman et al. n=6 healthy and n=6 AD subjects (median percentages are 44% and 19.6% in healthy and AD subjects, respectively, p = 0.065).

We further used this approach to analyze another, smaller, dataset of brain cells taken from 6 healthy and 6 AD subjects [9]. Similar to the first dataset, the number of reads from each cell was also highly variable ranging from ~300 to ~2,800 reads per cell. Focusing again on neurons, we found a similar effect of increased heterogeneity in AD neurons compared to healthy subjects (Figure 1C and S1G).

While we find evidence supporting our hypothesis in AD brains, this analysis strongly suggested that in order to rigorously test our theory, we must use an isogenic cellular system (iPSCs). In addition, it prompted us, instead of a droplet-based method with a large number of cells but relatively shallow sequencing depth per cell, to prefer the Smart-seq2 protocol with a relatively small number of cells, but which provides a superior sequencing depth over drop-seq methods and is more homogeneous in the number of reads obtained from each cell. This enabled us to use a more quantitative approach towards gene expression variability analysis.

To differentiate the iPSCs toward NPCs, we used a 20-day differentiation protocol [14] (Figure S2A-C and see Methods). Briefly, cells were dissociated into single cells, re-plated, and incubated in neural medium with the dual SMAD inhibitors SB431542 and LDN193189 and the Wnt signaling inhibitor XAV-939. From day 10 to 20 cells were treated with XAV-939 and an activator of Sonic hedgehog (Shh) signaling, SAG (Smoothened Agonist). At day 20, the cells become committed neuronal progenitor cells (NPCs) and express TUJ1 (Figure S2B).

In order to enrich for intact cells and filter out non-NPCs, we developed a fast and efficient protocol for cell harvesting, staining and sorting of NPCs into 96-well plates (Figure S2D and see Methods). In addition to sorting for live and non-apoptotic cells, we stained for PSA-NCAM, an outer membrane marker for NPCs. To minimize the processing time, we used a conjugated antibody for sorting. Typically, 30-70% of the intact cells were also positive for PSA-NCAM (Figure S3A-C). We then computationally filtered out cells based on the statistics of aligned reads, as well as the expression levels of neural and embryonic genes (Figure S3D-G), resulting in 40-90 cells for each cell line (Supplementary Tables S1 and S2).

### Genetic background dominates the transcriptional profile of cells over polyQ mutations

We first compared the single cell data to bulk data of RNA-Seq from 2000 cells from the same experiment taken in duplicates in the 72Q repeat isogenic system (Figure S4). Our analysis demonstrated a high correlation between the average single cell data and the bulk data (Figure S4A-D), in agreement with previous reports [15]. PCA analysis confirmed that WT and mutant cell populations were not composed of several distinct subpopulations, rejecting the possibility that changes between cell types are a result of different frequencies of subpopulations of cells (Figure S4E-G). We therefore used the average expression of the single cell data in order to compare between the different HD cell lines. We found that cell lines were grouped according to their genetic background, where isogenic pairs were more similar to one another than to HD-corrected cells from different backgrounds (Figure S4H). Therefore, the genetic background of the cells affects the cells’ transcriptome to a larger extent than the HD mutation itself (i.e., WT vs mutant), in agreement with other studies [11], highlighting the importance of using isogenic cell lines.

In addition, at the average expression level, we found that there are more differentially expressed (DE-) genes in the 180Q system compared to the 72Q system (Figure S4I). This is coherent with the idea that as the 180Q system contains a mutation that causes a more severe and earlier onset form of the disease compared to the 72Q system, more transcriptional changes are evident in early developmental stages in the 180Q system.

Among the several significantly and consistently DE genes in both the 72Q and 180Q systems, we found that three out of the five genes, namely *FN1, TGFB2* and *BHLHE40*, were previously shown to have altered expression either in the R6/2 HD mouse model or in hESC-derived NPCs and striatal-like post-mitotic neurons [11, 16–19]. In addition, consistent with our previous observation that isogenic pairs are more similar to each other compared to WT cells with different genetic backgrounds, we found that the number of DE-genes between mutant and WT non-isogenic pairs is larger than the number of DE-genes between mutant and WT cells within isogenic systems (Figure S4I).

### Neurological disorders show increased transcriptional heterogeneity

To test our hypothesis that the mutated NPCs have a larger cell-to-cell transcriptional heterogeneity, we used the intra-condition distance measure that was previously introduced [20]. In this paper, the authors compared between mouse ESCs grown in serum conditions, cells cultured with MEK and GSK3 signaling inhibitors (‘2i’), supporting ‘ground state’ conditions, and a KO cell line for the microprocessor complex subunit DGCR8, which is required for microRNA (miR) processing. Re-analyzing their raw data, we replicated their results demonstrating that 2i conditions, supporting a more homogeneous state [21], is indeed less heterogeneous compared to serum, while the DGCR8-KO cell population is the most heterogeneous (Figure 2A). We also used this measure for the AD datasets [8,9], confirming a significantly larger intra-condition distance between AD neurons and healthy subjects (p=0.042 for excitatory neurons in Mathys et al. and p=0.026 for neurons in Grubman et al., Wilcoxon signed rank test on the median correlations of patients). In our iPSC data, we found that while in the 72Q system the mutated and the isogenic cells are relatively similar (Figure 2B, blue graphs), the 180Q mutant cell population is evidently more heterogeneous compared to its corrected isogenic counterpart (Figure 2B, red graphs, mean difference 0.045, p ≪ 10^-100^, t-test). Interestingly, the ASD-associated mutant CHD8 cell line had the largest shift in this heterogeneity measure (Figure 2C, mean difference 0.15, p ≪ 10^-100^, t-test). These results are in line with our hypothesis and might further imply that the size of the effect is related to the stage of onset of the disorder.

**Figure 2.**
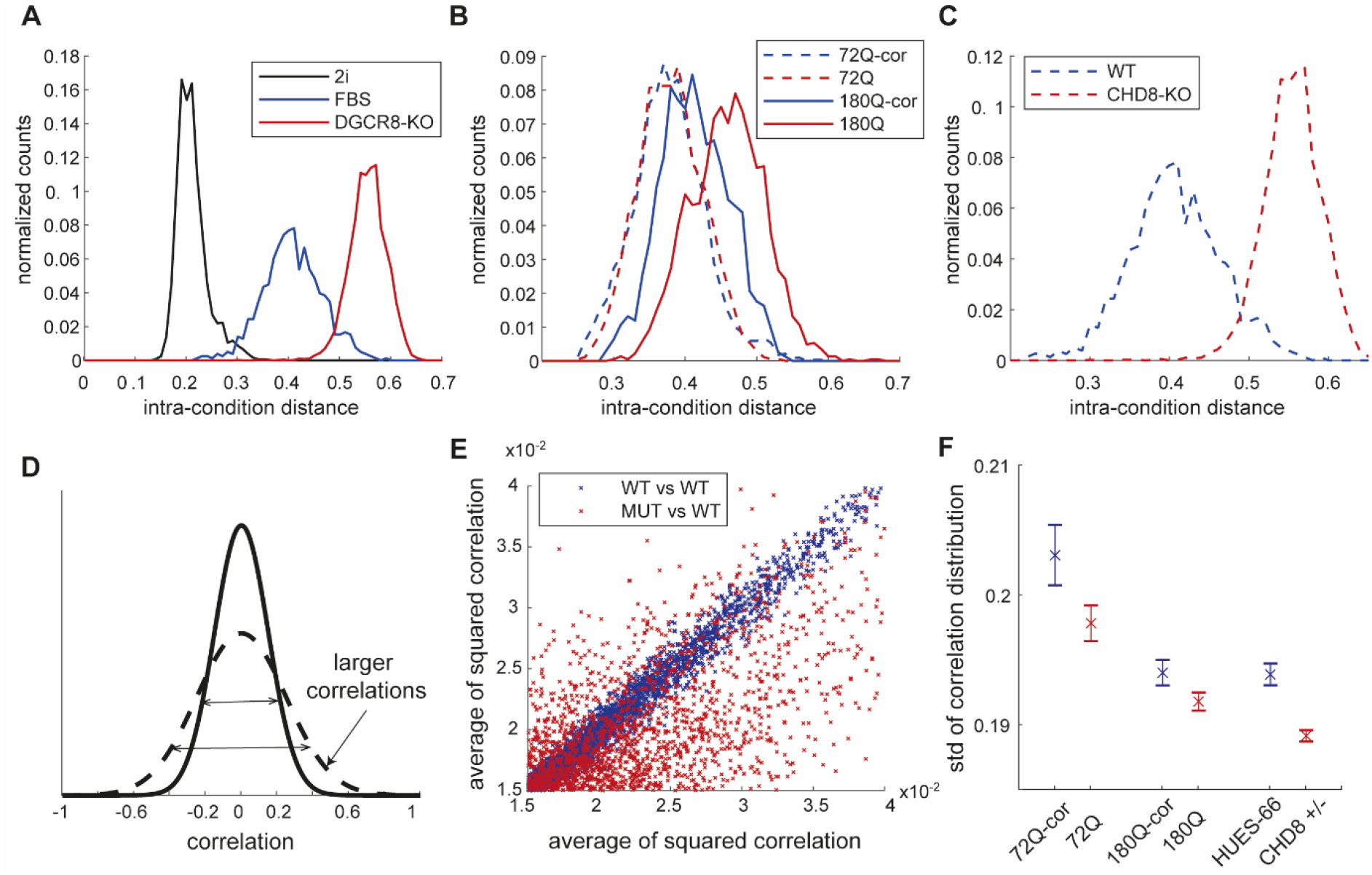
Cellular models of neurological disorders show increased transcriptional heterogeneity. (A) Heterogeneity of cell populations measured by ‘intra-condition distance’ (see Methods) in mouse ESCs comparing DGCR8-KO cells (red) and WT cells grown in either 2i (black) or serum (blue) media conditions. (B) Same as **a** for human NPCs derived from two HD isogenic pairs: mutant 72Q cells (dashed red) and their isogenic controls (dashed blue); and mutant 180Q (solid red) and their isogenic controls (solid blue). (C) Same as **a** for human NPCs derived from WT hESCs (blue) and CHD8^+/-^ isogenic cells (red). (D) A model for transcriptional disarray resulting in weaker correlations throughout the GRN. Shown is the distribution of correlations between all pairs of genes in the more organized state (dashed line) and the transcriptionally out-of-balance state (straight line). (E) The average of squared correlation of a gene with the rest of the genes. Each dot represents a single gene. Red, 180Q mutant vs corrected. Blue, two random pools of WT cells (see Methods). (F) Standard deviations of gene correlations in WT vs mutant cells in the 72Q, 180Q and CHD8^+/-^ isogenic systems. Error bars represents 1 std over 10 replicates (see Figure S4 and Supplemental Methods).

### The genetic regulatory network is globally less correlated in neurological disorders models

The intra-condition distance quantifies the overall dissimilarity (transcriptional dissimilarity in our case) between cells in the population. A complementary approach to the similarity between the different cells is the similarity between the transcriptional levels of the different genes. The expression levels of genes in the genetic regulatory network (GRN) of the cells may be either correlated, anti-correlated or neither (no correlation). Our hypothesis predicts that in the mutant state, the network would be less tightly controlled, and therefore the genes would be less coordinated. If this lack of coordination occurs at the transcriptional level, we expect to find smaller correlations (in absolute values) in the mutant state (Figure 2D).

To test this in our HD NPCs, we measured the correlation between each and every pair of genes in the network in the mutant and isogenic controls. Supporting our hypothesis, we found that both the HD and the CHD8 mutant cells show a significant (p-value < 10^-17^, 10^-14^, 10^-19^ for 180Q, 72Q, and CHD8 mutant systems, respectively, two-tailed t-test) decrease in gene correlations compared to WT (Figure 2F and S4J-L, and see Methods for details), with most of the genes showing a reduced average squared correlation with the rest of the genes (Figure 2E). A closer examination of this decrease in correlation revealed, in all systems, the presence of a cluster of several hundred (500-900) genes, which decrease together in pairwise correlations (Figure S4M-P). These clusters contained many ribosomal subunits (hypergeometric test FDR adjusted p-value ≪ 10^-10^), suggesting a global transcriptional disarray in the mutant cells.

### Variability comparison by modeling gene expression using a mixture model

In order to find genes that have increased variability in the mutant cell lines compared to the WT cells or vice versa, we modeled the gene expression distribution. Most previous studies that quantified gene expression variability from scRNA-Seq data, did not compare variability between two conditions but rather found the most variable (/stable) genes in a given experiment [20,22]. In most cases, the authors used the coefficient of variation (CV) measure as the basic statistic to compare variability and did not take into account the specific shape of the distribution. Furthermore, most studies performed the analysis in a group basis rather than a gene-by-gene basis [23]. In other works, mRNA expression distribution has been modeled using the bursty transcriptional model [4,24]. In this model, transcription is not a continuous process. Instead, the promoter of a gene can enter an active state during which mRNA molecules are synthesized until the promoter stochastically switches back to the inactive state. This model gives rise to a complex probability density function for the mRNA count in the different cells [25]. Under certain conditions, this theoretical distribution can be approximated by a negative binomial (NB) distribution [26,27]. Although the negative binomial distribution has been successfully applied for both single molecule fluorescent *in situ* hybridization (smFISH) [28] and scRNA-Seq data [26,29], genes which are not highly expressed in all cells are more difficult to model using only one NB distribution. Rather, it has been suggested to model the mRNA distribution using a mixture model with more than one distribution [28].

Empirically, we found that in our data, most of the genes faithfully corresponded to either a Gaussian distribution, an exponential distribution, a uniform distribution around 0 or a mixture of those (Figure 4A-E and S5A-C). This model also has computational advantages, as the NB model has no closed formula for the maximum-likelihood solution, which is even more difficult when a mixture-model parameters are to be deduced.

**Figure 3.**
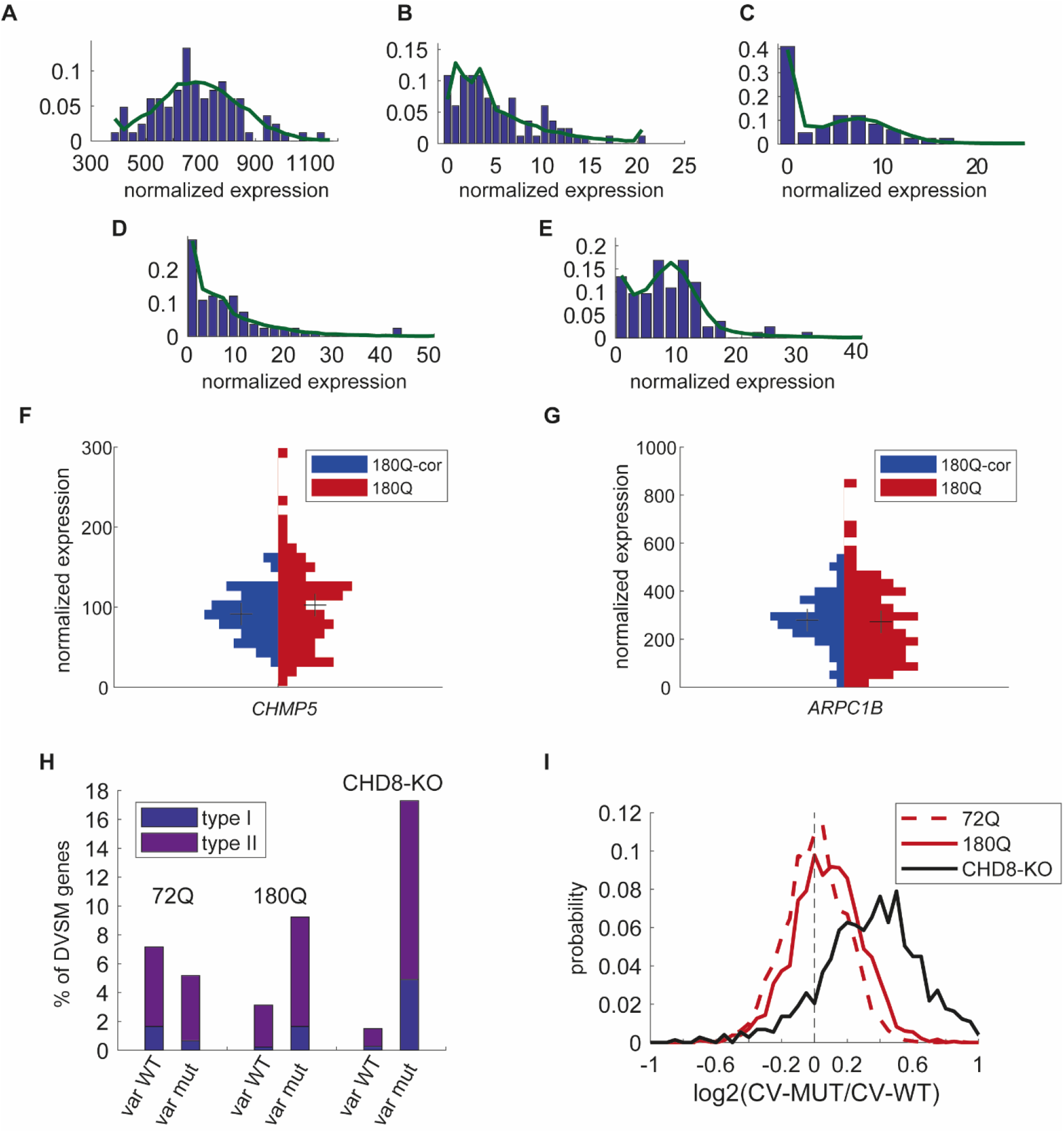
Models of neuronal diseases show a global increase in transcriptional variability. Examples of best mixture model fit (see Methods for details) of gene expression distribution. (A) Pure Gaussian distribution. (B) Dominant Exponential distribution. (C) A mixture of Exponential and ‘Uniform around 0’ distributions. (D) A mixture of Exponential and Gaussian distributions. (E) A mixture of Gaussian and a Uniform around 0 (zero-spike) distributions. Green line marks the probability density function of the best fit. (F-G) *CHMP5* and *ARPC1B* expression is type I and type II more variable in 180Q compared to 180Q-corrected cells, respectively (see Methods for details). Shown are the histogram of the normalized expression values. Black crosses represent the mean expression. (H) The number of genes which have a similar mean expression in both WT and mutant cells are differentially variable (for details see Methods) in 72Q, 180Q and CHD8^+/-^ isogenic systems. (I) The distribution of the log ratio of the CV between mutant and WT for the three isogenic systems. 72Q: dashed red line, 180Q: red solid line, CHD8^+/-^: black solid line.

**Figure 4.**
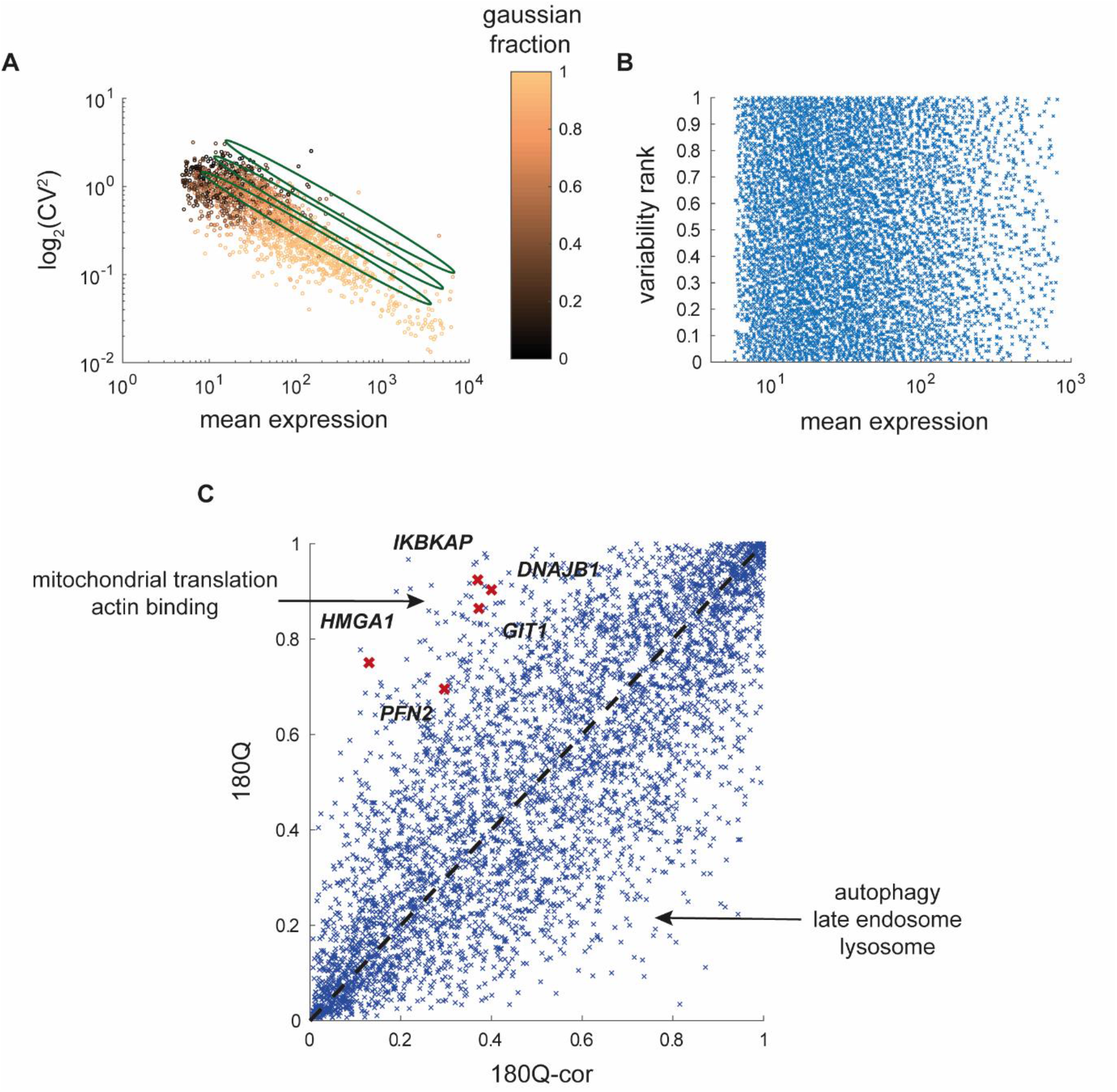
Functional analysis of differentially variable (DV-) genes. (A-B) The mean expression is correlated with the Gaussian fraction of the mixture model fit and reversely correlated with the trimmed-CV. Each dot represents one gene. Genes are color-coded based on the Gaussian component in the mixture mode fit where orange represents pure Gaussian distribution and black represents no Gaussian fraction. The CV is transformed in a layered fashion (green ellipses) into the variability ranked score (B) which is independent of the mean. (C) Shown is the variability rank score of all genes in the 180Q mutant vs WT cells. Genes which are known to interact with the HTT protein and are significantly differentially-variable are marked in red.

As we were interested in isolating the variability-related effects from the average expression difference, we first limited our analysis to genes, which have a similar average expression in both WT and mutant cells (see Methods for details). Our model allowed us to determine in which of two conditions a given gene is more variable. First, if the expression distributions in both WT and mutant cells could be dominantly fitted by Gaussian distributions and the standard deviation in one of the conditions was significantly larger, then the gene was considered as more variable in this condition (type I, see Methods). Second, if the expression distribution of a gene could be dominantly fitted by a Gaussian distribution in only one of the conditions, this means that in the second condition there are more extreme values (zeros and very large expression levels), and therefore it was considered more variable in the second condition (type II, see Methods). Examples include *CHMP5* (type I, std ratio of 1.46-2.29 with 95% confidence interval, Figure 3F) and *ARPC1B* (type II, Gaussian fraction of 0.98 in WT vs 0.49 in mutant cells, Figure 3G), where the mutant cells (red) show a wider normalized expression distribution than their isogenic counterparts (blue, see Methods for details).

### Neurological disorders models have larger gene expression variability

Using our modeling approach, we were able to compare the expression variability changes between isogenic pairs of WT and mutant cells for genes with similar average expression in both conditions (Figure 3H). In addition, we also used the CV as a complementary approach (Figure S5E-F and Methods). We hypothesized that because the mutant cells are less tightly regulated, as a result of the mutation, they will have more genes with increased variability compared to their WT isogenic counterparts. This effect is expected to be stronger in the 180Q juvenile onset cells compared with the 72Q cells, and even stronger in the CHD8 mutant cells. Reassuringly, we observed that while in the 72Q cells the mutant and WT cells are relatively similar, with slightly more genes which show increased variability in WT cells and a weak CV bias towards the mutant cells (p = 0.0591, Wilcoxon signed rank test, and Figure 3H, left bars), in the 180Q system there is a significant CV bias toward the mutant cells (p < 10^-50^), which also show far more genes with larger variability (Figure 3H, middle bars). This effect is further enhanced in the CHD8 mutant cells (p < 10^-90^, and Figure 3H, right bars). Overall, in agreement with our hypothesis, the difference in the number of genes, which are more variable in the mutant cells compared to WT, increases as the reference age of onset of the neuronal disorder decreases.

To assure that our results are not an artifact of a possible whole genome duplication, we performed e-karyotyping analysis for all cell lines (inferCNV of the Trinity CTAT Project. https://github.com/broadinstitute/inferCNV, data not shown). We found that all HD-related cells as well as the CHD8^+/-^ cell line showed no signs of chromosomal aberrations. Unexpectedly though, the HUES-66 WT cell line showed a signature of chromosomal duplication in chromosomes 12 and 20. While we would not expect such a genomic aberration to result in a global decrease in expression heterogeneity, we nonetheless repeated the analysis for the ASD isogenic system removing all the genes within the suspicious chromosomal regions. Although the removal of the suspicious chromosomal regions from the analysis does not completely rule out the possibility of karyotype-related side-effects, it does control for the major side-effect arising from the bias in the estimation of the normalized expression levels of all genes. The CHD8^+/-^ cells still showed a highly significant CV bias (Figure S4D), suggesting, as expected, that the chromosomal aberration in the HUES-66 WT line, if present, is not the cause for the significantly increased expression variation observed in the CHD8^+/-^ line.

In order to understand which functional pathways in the cell are affected, we ran a gene annotation analysis for the ‘differentially variable with similar mean’ (DVSM) genes in the 180Q system. We found no prominent characteristics for genes that were more variable in the corrected cells. By contrast, the genes that were more variable in the mutant cells were enriched for cellular energy production related terms (Table S3), such as tricarboxylic acid cycle (hypergeometric test FDR adjusted p-value = 1.21×10^-04^), NADH dehydrogenase complex assembly (p = 1.59×10^-6^), and aerobic respiration (p = 6.51×10^-05^). Interestingly, mitochondrial damage has been implicated in HD at the mitochondrial DNA level [30] and the mRNA level [31], as well as at the level of enzymatic activity [32–34] and cell survival [30] in both mouse and human. In addition, actin polymerization related genes, including several subunits of the actin related protein 2/3 complex were more variable in the mutant cells. Among these genes was Profilin, which directly interacts with HTT, and was shown to inversely correlate with aggregate formation in polyQ cellular models as well as disease progression in patients [35, 36]. These results suggest that altered transcriptional heterogeneity is not random and that transcriptional variability is a regulated process, related to the cell’s state.

### Extending the analysis for all genes reveals differentially variable (DV) genes in HD

The analysis thus far concentrated on the genes for which we could isolate the variability effect from the average expression level changes, by using only genes that have similar mean expression in WT and mutant cells. However, there are many other genes for which both the average expression and the variability are different in the two conditions. This comparison is not straightforward, however, as the CV is reversely correlated with the mean (Figure 4A). This bias has been previously observed both at the protein level [37] and the mRNA level [22]. The bias also holds for the mixture model approach, as the fraction of the Gaussian part of the fit is correlated with the mean as well (Figure 4A).

In order to correct for the CV-mean bias, previous studies have either statistically modeled gene expression using a Bayesian approach [38] or used the trendline of this correlation [23, 39]. We used an approach similar to the latter to correct for this dependence (see Methods). For each condition, we ranked the genes based on their CV relative to that of the genes with similar mean. This resulted in a variability rank measure, which is uncorrelated with the mean expression (Figure 4B).

In general, most of the genes had similar ranked-variability in both WT and mutant cell lines (Figure 4C). Expectedly, many of the genes that had lower ranked-variability were metabolic and splicing and translation-related housekeeping genes. Genes which were more variable in mutant cells, were enriched for mitochondrial ribosomal subunits (Table S3). In addition, genes which are known to interact with the HTT protein itself, were enriched in the most differentially variable genes that were more variable in the mutant cells (Figure 4C, red crosses, p = 0.0087, hyper-geometric test). By contrast, genes that were more stable in the 180Q mutant cells were enriched for autophagy, lysosome and endosomal-related processes (Table S3), consistent with the idea that the mutant cells must tightly control the cellular mechanisms that are responsible for damage control as a result of the mutation.

Searching for a cellular mechanisms that could mediate the changes in gene expression variability between the WT and the mutant cell lines, we ran a motif analysis to search for a binding site that is enriched for the differentially variable genes [40]. We found that while the DV genes, which were more variable in the HD mutants, are enriched for NKX2-1 motif (Figure S5G), the genes which are more variable in the WT cells are enriched for the glucocorticoid related GMEB1 motif (Figure S5H). This might suggest that an upstream regulator is, at least partly, responsible for the changes we observe in expression variability.

### Phenotypic consequences of disrupting differentially variable genes in HD

Our analysis revealed that the HD mutation results in changes in gene expression variability. But are these changes relevant to the pathology of the disease? In order to test this, we used a reporter system of human embryonic kidney 293 (HEK) cells that we developed (Figure 5A). These HEK cells were stably infected with a GFP reporter fused to the 5’ region of the human *HTT* gene containing a polyQ-encoding stretch of 105 repeats. The GFP-105Q proteins can form intracellular aggregates as in HD, although not all cells that express the GFP-105Q plasmid form aggregates.

**Figure 5.**
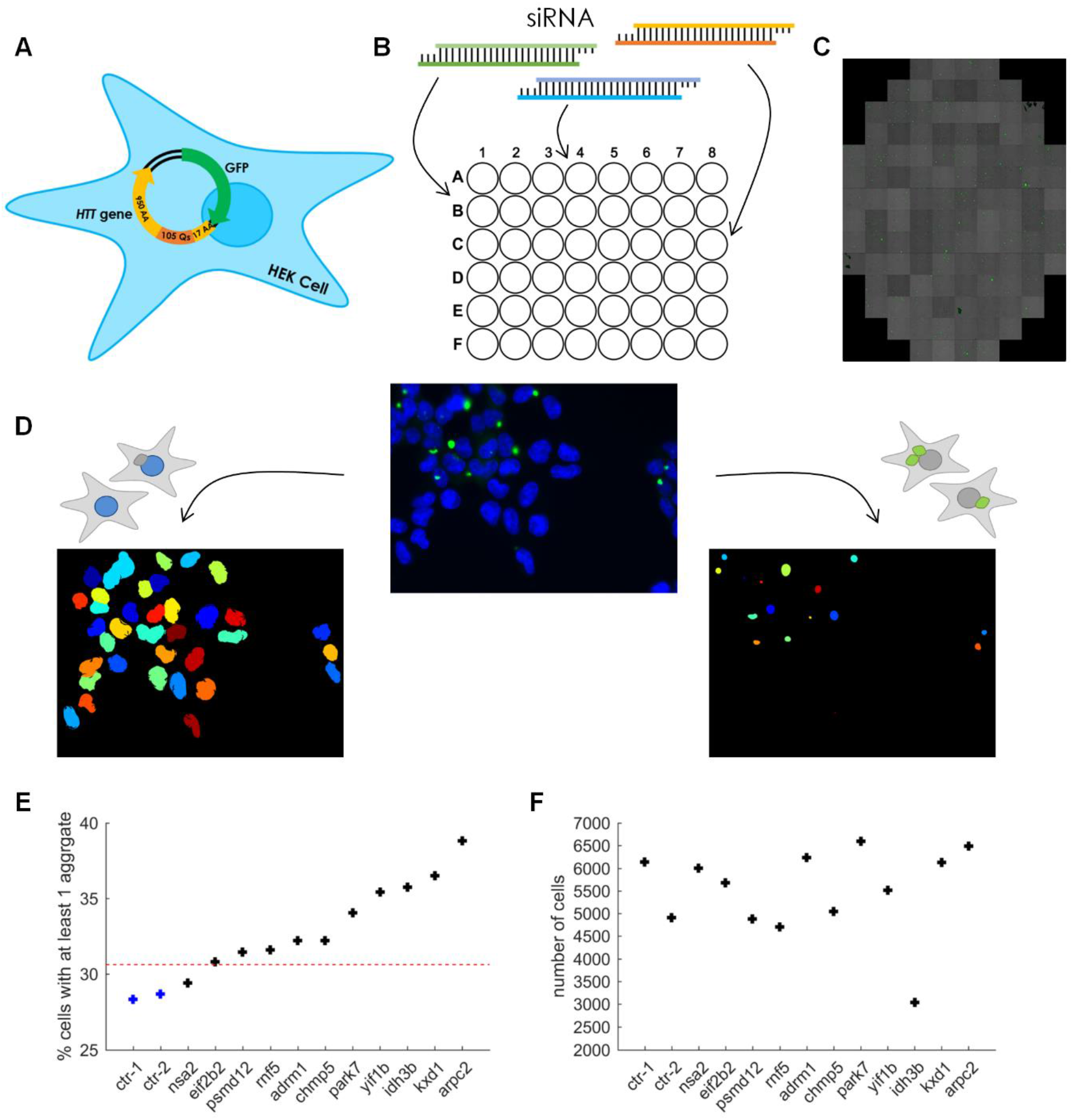
Functional validation assay for HD potentially-related genes. (A) Schematic representation of HEK293 cells and 180Q NPC infected with a GFP-105Q construct. (B) A heterogeneous population of GFP-105Q cells with 50% of the cells showing aggregates were transfected with siRNAs to knockdown a specific target gene (See Methods). (C) After 48 (for HEK) or 96 (for NPC) hours, cells were scanned using an automated fluorescent microscope (high content imaging). Aggregates are in green. (D) Image processing to identify nuclei (bottom left) and aggregates (bottom right). Original images containing both Hoechst-stained nuclei (blue) and polyQ aggregates (green) were segmented separately. (E) Percentage HEK cells, which have at least one polyQ aggregate, following knockdown of different target genes, listed below. Red dashed line represents the average of 3 standard deviations from the average of non-treated (WT) and scrambled (SC) samples. (F) Total number of nuclei for each target gene siRNA assay. (G-H) Same as E-F for NPCs. Red boxes represent DV genes with increased variability in mutant cells, orange boxes represent non-DV genes previously associated with aggregates and black boxes represent control genes. Green dashed line represents the average of 2 standard errors of the mean from the average WT and scrambled samples.

To test the functional importance of differential gene expression variability, we designed an siRNA screen where the HEK cells were transfected with specific siRNAs to target genes that we found to be differentially variable in our HD cells (Figure 5B). We chose genes that had the largest effect in both 180Q and 72Q systems. We mixed GFP-105Q cells that expressed the reporter in 100% of the cells with cells that did not express the reporter at 1:1 ratio to achieve an initial pool of cells with 50% of the cells expressing the aggregate reporter. 48 hrs after the addition of the siRNA, we stained the cells with Hoechst to mark the nuclei and used an automated fluorescent microscope to generate images from more than 100 fields per gene (Figure 5C-D).

To analyze the images, we used a semi-automated approach. First, in order to automatically recognize the nuclei and the aggregates in each image, we built a protocol in CellProfiler program [41] and fine-tuned the parameters based on a small fraction of the images (Figure 5D). We used permissive parameters to identify most of the potential objects (Figure S6). This resulted in several thousand image-processed cells for every gene. We estimate that our pipeline recognizes 80-90% of the cells and nuclei with little or no false positives.

Using our pipeline, we could efficiently evaluate the effect of the knockdown of the different genes on the polyQ aggregates. We compared the fraction of cells with aggregates in each knockdown to untreated cells and to scrambled controls. We found that knockdown of DV and DVSM genes that were more variable in both the 180Q and the 72Q isogenic systems (*KXD1, IDH3B, YIF1B* and *RNF5*) or only in the 180Q system (*ARPC2, CHMP5, PARK7, ADRM1*, and *PSMD12*) had the strongest increase in aggregate formation (Figure 5E). This was in contrast to *NSA2*, which is DV in the 180Q system and to our control gene, *EIF2b2*.

Overall, for the great majority of the DV genes we tested, the knockdown resulted in a significant increase in the fraction of cells with aggregates (Figure 5E). The fraction of cells with aggregates was not correlated to the number of detected cells in each condition, suggesting that these results are not a result of the potentially slower cell cycle progression in the treated cells (Figure 5F). While these results nicely demonstrate that the differentially variable genes are potentially associated with HD pathology, the HEK293 cell line is less relevant to HD and is karyotypically abnormal. We therefore developed another polyQ aggregate reporter system, introducing the same 105Q-GFP cassette into our differentiated 180Q NPCs. Repeating the analysis using an NPC calibrated pipeline, we again evaluated the effect of knockdown of 14 DV target genes as well as 15 control genes. We chose control genes which show similar average expression in WT and mutant cells but are not DV. Among them, we included genes *RBM39*, *CCT5* and *CCT7*, which are not DV but which were previously implicated in aggregate formation [42–44]. Among them is also *COMMD6*, which shows a similar expression in WT and mutant cells, but is more variable in WT cells. When we knocked down these genes and automatically scored aggregates, we found that nine out of 12 control genes showed no effect on aggregate formation (Figure 5G, black bars). WT cells and NPCs treated with scrambled siRNAs also showed no response, as expected (Figure 5G, blue bars). In contrast, we found that 10 out of 14 DV target genes, showed a significant and larger increase in polyQ aggregate formation in the NPCs (Figure 5G, red bars). This was also the case for 2 out of the 3 genes which were not DV but which were previously associated with aggregate formation: *RBM39* and *CCT7* (Figure 5G, orange bars). The most profound effect was observed for *PARK7* (a.k.a. DJ-1), which was shown to be heavily implicated in aggregate formation in both Parkinson’s and Huntington’s diseases [45]. Overall, these data suggest that the high-confidence differentially variable genes we identified are relevant for polyQ formation and disease pathology.

## Discussion

Many neurological disorders tend to manifest many years after birth, although the inherent cause in many cases, such as the mutant polyQ protein in HD, is already present in the germline. Here we aimed to understand to what extent HD, and possibly other neurological disorders, is a result of an accumulative deterministic process or caused by increased imbalance at the transcriptional level.

The commonly accepted model is the accumulative deterministic model; The mutant proteins, or other types of damage, gradually accumulate in the cells where they induce toxic effects leading to cell death [46]. The deterministic model predicts that as time goes by, differences between the WT and mutant cells accumulate, and these can be easily detected when averaging over cell populations. Supporting this model, most HD models, for example, exhibit many transcriptional changes compared with WT [16,17,47].

In our work, we began by analyzing existing scRNA-Seq of AD brains and found increased transcriptional heterogeneity in neurons in general, and excitatory neurons in particular. However, these cells displayed a very wide variation in sequencing depth, and therefore in most cases and most cell types, experimental noise prevented thorough analyses. Here we took advantage of two juvenile HD iPSCs, containing either 180Q and 72Q, together with their isogenic controls, and generated NPCs via *in vitro* differentiation. Since polyQ length is negatively correlated with the age of onset, the 180Q cells are expected to manifest greater phenotypes than the 72Q cells, enabling us to directly test our hypothesis. We found that in line with the deterministic model, there were significantly more DE-genes in the 180Q system compared to the 72Q system.

By contrast, measurements in HD patients showed that the probability of cell death remains the same during the course of the disease [48]. This is consistent with the stochastic model, which predicts that because the changes between HD and WT are at the stability levels rather than the absolute levels of the phenotype, changes will be noticeable only at the single cell level. As most experiments compare only cell populations in bulk, it is currently still hard to find more evidence for this model.

Our results suggest that the HD and CHD8^+/-^ mutations lead to differences in expression variability of many genes. The potential cellular mechanisms that could mediate these changes are at the level of mRNA synthesis (i.e., transcription) or at the level of mRNA stability. At the level of mRNA synthesis, these changes can be mediated by changes in the binding of transcription factors (TFs) to regulatory elements of the genes, as well as changes in post-translational histone modifications (HMs). Indeed, previous studies suggested that both types of changes are linked to variability at the mRNA level [39,49,50]. As CHD8 is a DNA helicase that acts as a chromatin remodeler, its loss of function could therefore alter the dynamics of mRNA production in many genes and increase the noise levels. In HD, the mutant protein localizes to the nucleus and binds many TFs. We found that several of the proteins that physically interact with HTT are more variable in the mutant cells compared to WT, including HMGA1 and IKBKAP, both of which directly bind chromatin. This suggests the intriguing possibility that the impaired interaction of mHTT with these proteins may result in a global disarray caused by changes in gene expression dynamics of genes that are affected by these proteins and are otherwise robustly expressed in WT cells.

In addition to these direct effects, the presence of mHTT can also produce indirect effects downstream of its impaired interaction with its different binding partners. Supporting this idea, searching for binding motifs in the DV-genes revealed that while the genes which were more variable in the HD mutants, were enriched for NKX2-1 motif, the genes which were more variable in the WT cells, were enriched for GMEB1 motif. While these factors are not directly regulated by HTT, they are both linked to HD. GMEB1 binds to glucocorticoid modulatory elements and enhances their sensitivity to glucocorticoids. Glucocorticoids have been previously shown to contribute to HD pathogenesis in an HD Drosophila model [51]. Interestingly, mutations in the *NKX2-1* gene, which encodes for a protein that regulates the expression of thyroid-specific genes, cause benign hereditary chorea, a characteristic symptom of HD. Therefore, our results raise the possibility that the changes in gene expression variability in the disease state are mediated, at least to some extent, by DNA regulatory elements, which can recruit known TFs. Future research would reveal disease-related changes in the binding dynamics of different TFs.

Our analysis showed that suppressing genes that have variable expression in mutant vs. corrected cells, causes, in most cases, elevated polyQ aggregation, a pathology associated with HD. This suggest that the DV genes we identify in this type of analysis are, at least in most cases, relevant for disease pathology. Among the control genes that had a significant effect in NPCs were *RBM39*, which was previously associated with cellular aggregates in yeast [42], as well as *MRPL17*, encoding for a mitochondrial protein, and the Spinocerebellar Ataxia 10 associated, tRNA-processing gene, *DTD1*. In addition, inhibition of ECM1, which is associated with lipoid proteinosis that may result in epilepsy, neuropsychiatric disorders, and spontaneous CNS hemorrhage, resulted in a significant effect [52]. Interestingly, knockdown of HDAC3, which has a similar average expression and variability in HD NPCs, and was previously shown to be related to HD cognitive pathology and CAG repeat expansion but not to aggregate formation in mouse HD models [7], showed a slightly significant increase in cellular aggregates. Finally, although both CCT5 and CCT7 are known to contribute to protein aggregation through autophagy regulation [44], only CCT7, which was shown to interact and affect GPCR aggregation [43], showed an effect in our NPCs.

It should be noted that by knocking down the genes we did not directly modify the variability in their expression levels but only their expression level. How can variability be directly manipulated for a specific gene without affecting other genes as well? One approach can be based on the DAmP multiple copy array (DaMCA) method that was developed in yeast [53]. The idea behind this method is that destabilizing the relevant mRNA such that it has a short half life time, while sufficiently increasing the number of copies of the gene (from the 1-2 copies) can result in the same average number of mRNA molecules as the original cells, but with smaller variability in the number of mRNA molecules. In addition, the variability in mRNA levels can be reduced by incorporating gene regulation motifs that are expected to reduce noise levels. The simplest motif that can decrease noise levels is the negative feedback loop [54]. Thus, if the inserted genes also include a regulatory element that binds a repressor encoded by the gene, noise levels can be further reduced. The introduction of these types of constructs into our system may be challenging, because of the need to titer the number of gene copies in order to maintain the same average expression levels in the NPCs rather than at the pluripotent undifferentiated state. However, recent advances in genomic editing techniques in mammalian cells make it more tangible.

Finally, since our model suggests that increased transcriptional heterogeneity occurs much earlier than the time of disease onset, it may have a diagnostic value. While it is impossible to measure transcriptional variability in neuronal cells *in vivo*, iPSCs allow the derivation of patient neurons in culture. It should be noted, however, that in order to properly estimate transcriptional variability, isogenic baseline controls are required, which are impossible to obtain for non-genetic diseases. Another potential baseline is using the patient own cells derived at different ages, either by directly trans-differentiating the patient’s cells to neurons [55], or by aging the cells in culture, e.g. using Progerin [56,57]. Further research will be needed in order to estimate the interplay between the effects of age and different NDs on transcriptional variability.

## Conclusions

In our work, we showed that human iPSC-derived NPCs exhibit an increase in transcriptional disarray. We demonstrated this transcriptional imbalance with three different approaches. Most importantly, we found that in the 180Q isogenic system a large number of genes are differentially variable with the same mean expression in WT and HD and are more variable in HD. Overall, our work suggests that gene expression heterogeneity can contribute to HD and possibly other genetic disorders. Therefore, treatments that aim at stabilizing the transcriptional state in the diseased cells, possibly through changes in binding of TFs and HMs, may be beneficial.

## Methods

### Cell Lines

The following cell lines: the 180Q isogenic system: 180CAG, 96-ex (180CAG-corrected), and the 72Q isogenic system: HD-iPS-72Q, (C116 HD-iPS-72Q-corrected) were generated by M. Pouladi and L. Ellerby, respectively. Full experimental details can be found in [11] for the 180Q isogenic system and in [10] for the 72Q isogenic system. The CHD8 isogenic human embryonic stem cell lines were HUES66 [58] for wild-type and AC2 for the heterozygous loss-of-function mutant, created using CRISPR/Cas9 mutagenesis (Shi X et al., Submitted).

### Cell culture and neuronal differentiation

hESCs, and hiPSCs were cultured on mitomycin-C treated MEF feeder layer in standard ESC media (DMEM containing 15% ESC-qualified fetal bovine serum (FBS), 0.1 mM nonessential amino acids, 1 mM sodium pyruvate, 2 mM L-glutamine, 50 μg/ml penicillin-streptomycin, 100 μM β-mercaptoethanol, 8 ng/ml bFGF). For neuronal induction [14], hESC and hiPSC colonies were separated using accutase and passed through 70 μm cell strainer followed by 2 passages of 25 minutes to dispose of MEF cells, and then seeded on Matrigel as single cells in hESC conditioned media, which was replaced the next day with human neural induction media (DMEM HAM’s F12, with 50 μg/ml penicillin-streptomycin, 2 mM L-glutamine, 1:100 of N2 (Invitrogen, cat. 17502-048), 1:50 of B27 (Invitrogen, cat. 12587-010)), supplemented with 2 SMADi (20 μM SB, 100 nM LDN) and 1 μM XAV-939 for 10 days. From day 10 to 20 of differentiation, cells were incubated with neural induction media supplemented with 50 nM SAG and 1 μM XAV-939, inducing the cells to become neural progenitor cells (NPCs). NPCs were cultured for up to 5 passages on Matrigel with neural induction media supplemented with 40 ng/ml bFGF (Peprotech: (AF)100-18B), 40 ng/ml hEGF (Peprotech: (AF)100-15), and 7500 units hLIF (Sigma-Aldrich: L5283). The HD 72Q and 180Q isogenic systems were differentiated separately (i.e. as biological repeats) and were harvested on the same day. The ASD-related CHD8^+/-^ hESCs (CRISPR-mutated and isogenic controls) were differentiated and harvested separately from the HD systems.

### FACS

In order to prepare the cells for FACS sorting, medium was removed and cells were washed once with Dulbecco’s Phosphate Buffered Saline (D-PBS) and incubated with TrypLE™ Select Enzyme (Thermo Fisher scientific, cat. 12563029) solution (300 μl/well) at 37°C for 1 minute until cells were dissociated from the bottom of the culture dish. TrypLE was then counteracted with D-PBS with 10% FBS and cells were filtered through dry 70 μm mesh/cell strainer and counted. Cells were pelleted at 1200 RPM (300 g) for 10 min, and re-suspended in Binding buffer from the cell apoptosis kit (MBL, code: 4700). AnnexinV-FITC, PI (MBL, code: 4700) and ANTI-PSA-NCAM conjugated-Ab (Miltenyi Biotec, cat. 130-093-273) were added according to the manufacturer’s instructions. Cells were incubated in 4°C in the dark for 30 minutes. The reaction solution was then diluted in 5 ml PBS with 10% FBS and cells were pelleted and re-suspended in PBS with 1% FBS. Cell suspensions were then filtered again through dry 70 μm mesh before acquisition on a flow cytometer. For FITC and PI staining, cells with no staining served as controls. For PSA-NCAM staining, hiPSC were used as negative controls. Cells were sorted using 100 μm nozzle into a 96-well plate (Eppendorf, cat. 951020401) one cell in a well. Wells contained 5 μl/well TCL-buffer (QIAGEN, cat. 1031576) with 1% 2-Mercaptoethanol. Bulk controls were sorted into 75 μl DNA/RNA Shield (Zymo Research, cat. R1100-50). Following sorting, plate was immediately frozen on dry ice and kept at −80°C.

### Immunofluorescence and immunohistochemistry

For Immunofluorescence (IF), cells were fixed in 4% Paraformaldehyde (PFA), permeabilized with 0.5% Triton X-100 and blocked with 5% fetal bovine serum (FBS). Appropriate Alexa 488-conjugated secondary antibodies was diluted 1:500 and mixed with DAPI. Images were acquired using an Olympus IX71 X-cite epifluorescence microscope. The following primary antibody was used: Anti-beta III tubulin (ab18207) diluted 1:250. The secondary antibody was goat anti-rabbit IgG conjugated with Alexa-488 (Invitrogen).

### scRNA-seq protocol and pre-processing of scRNA-seq data

The scRNA-seq was generated from frozen dorsolateral prefrontal cortex from participants in the Religious Orders Study or Rush Memory and Aging Project (ROSMAP) [59].

For scRNA-seq generated in this work, we used a modified Smart-seq2 protocol as described [12,13]. We sequenced the libraries with a NextSeq 500 (Illumina). Reads were aligned to the human RefSeq reference genome (GRCh38) using bowtie2 v 2.2.5 [60]. We used RSEM version 1.2.31 [61] to quantify gene expression levels for all RefSeq genes in all samples. We further normalized the output transcripts per million (TPM) values based on quantile normalization [62] to remove bias that results from a small number of highly expressed genes.

### Bulk RNA-seq

We extracted total RNA using the Quick-RNA Mini Prep (Zymo Research) and followed the vendor protocol with a DNase treatment step. We prepared the RNA-seq libraries using the original Smart-seq2 protocol [12] from 0.5 ng total RNA input of each. We sequenced ~1 million 50-bp paired-end reads per sample on an Illumina NextSeq 500 sequencer.

### siRNA assay

After running the PulSA method [63] in flow cytometry, HEK-105Q cells or 180Q-105Q NPCs with and without aggregates were collected. 30,000 cells were plated in 24 well plates, such that 50% of the cells are from the aggregates containing fraction and 50% of the cells are from the fraction with no aggregates. Final concentrations of 25 nM of siRNA pools, including a scramble control, were used in transfections. Transfection was conducted 24 hours after plating (using Mirus reagent). Media was changed after two days. Scanning using Hermes WiScan was conducted 3 days after transfection. Hoechst was used to stain nuclei (1 μg/ml). Primers used for knockdown validations are shown in Table 1.

**Table 1.**
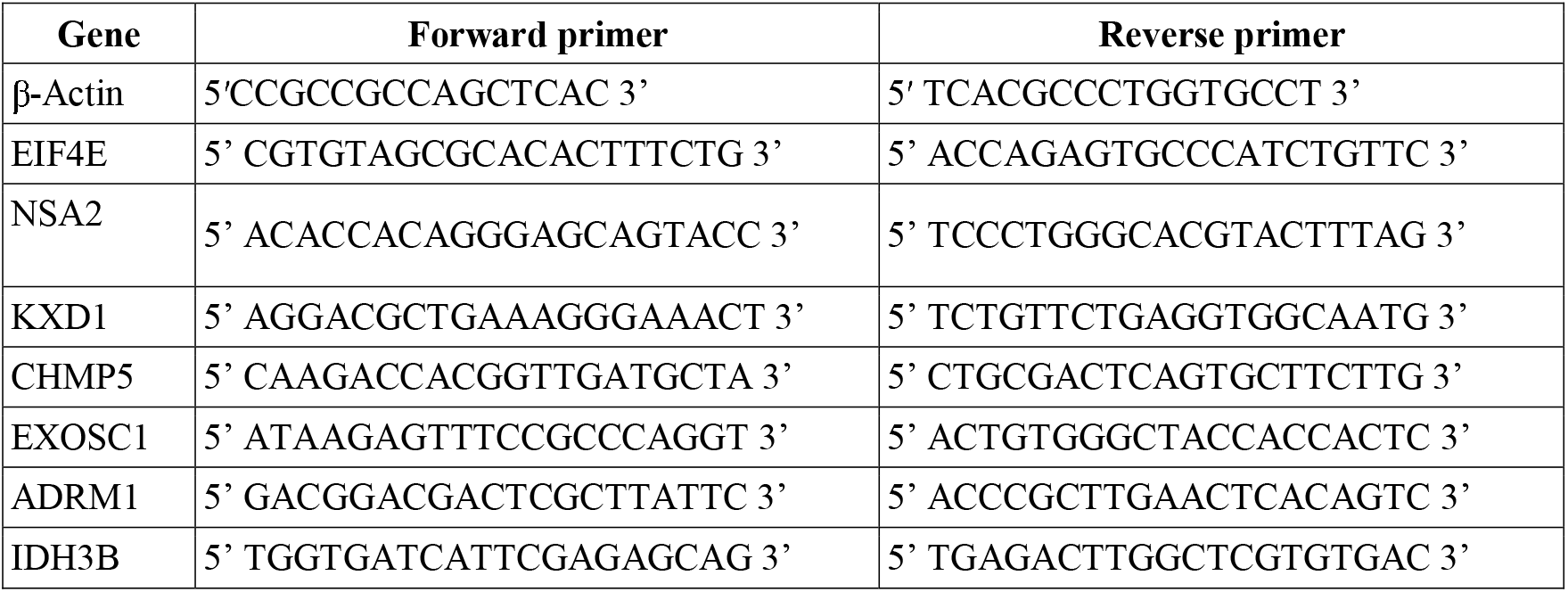
Primers used in real-time PCR.

### scRNA-seq data quality control

Prior to analysis, cells were filtered based on QC criteria. First, cells with less than 2×10^5^ aligned reads or less than 15% of all RefSeq genes detected were filtered out. For the NPC experiments, in order to eliminate non-NPC cells, we defined the neural-to-embryonic (N-E) index as the sum of expression of the NPC markers *NES* and *TUBB3* divided by the sum of expression of the two ESC markers *POU5F1* and *NANOG*. Cells that had an N-E index smaller than 2 were discarded from further analysis. Finally, to remove extreme outliers from the data, we calculated for each experimental condition the Spearman correlation between every two cells. We then used the 5-MAD criterion to remove cells with a very low average correlation with the rest of the cells (i.e., cells with average correlation with the rest of the cells that was more than 5 MADs away from the median were filtered out). We repeated the QC with the Seurat pipeline [64] and obtained identical results to our two initial filtering steps (prior to the exclusion of suspected non-NPCs).

### General heterogeneity measure for Alzheimer’s disease-related data

For each of the 48 subjects from Mathys et al. [8] cells were divided into groups based on the classification to the main neuronal types used in the original study. Then, for each subject for a given cell type, genes were ranked according to their expression level in each individual cell, and the top 200 genes that had the largest average rank were used. Next, for each gene the fraction of cells that expressed that gene was calculated, and the median detection level across the 200 genes was used as the statistic that represents the level of heterogeneity for that cell type in this subject, where small values close to 0 correspond to high level of heterogeneity and 1 corresponds to low level of heterogeneity. The same trend was kept also when top 100 and 500 genes were used. For the smaller dataset from 12 subjects from Grubman et al. [9] we used the cells that were classified as neurons by the authors and used the top 100 genes.

### General heterogeneity measures for NPC data

Genes were considered differentially expressed if the sum of the average of log2(normalized counts+1) in the two samples were larger than 3 and they had at least a 2-fold difference with FDR adjusted (Benjamini-Hochberg) p-value < 0.05.

Intra-condition distances (Figure 2) were computed as previously described [20], as (1-Pearson correlation coefficients) between cells among all genes with expressed on at least 50% of the cells and average expression larger than 5 normalized counts in both conditions. For the AD data all genes were used and significance level was calculated based on Wilcoxon rank-sum test between the medians of the intra-condition distance among all cells in a subject.

Gene correlation comparison was based on spearman correlation between every pair of genes. We mark by 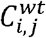 the spearman correlation between gene *i* and gene *j* in WT cells and in 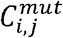 the correlation between gene in mutant cells. The average squared correlation of gene *i* is in condition *cond* is defined as 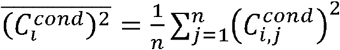 where *n* is the number of genes. The squared correlation difference between two genes *i,j* is defined as 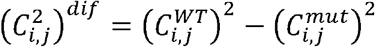. For more details see Supplementary Methods.

Clustering was conducted based on squared correlation differences for all pairs of genes. Empirically, we found that 2 clusters best describe all isogenic systems. We found the main cluster of genes that decreased in correlation between WT and mutant cells by running the k-means algorithm in Matlab using 2 clusters, 6 replicates and MaxIter=10000. The algorithm was run 10 times and the largest cluster was selected.

### Mixture model for gene expression data

To describe the shape of gene expression levels we used a mixture model of three distributions: A Gaussian, an Exponential distribution and a Uniform distribution between 0 and 1. The total number of parameters in this mixture model is 5, including 2 for the proportions of the three distributions, two for the normal distribution parameters and one for the exponential. In order to find the best fit for the expression level distribution of every gene, we used the Expectation-Maximization (EM) algorithm, which is based on the maximum likelihood criterion. For more details see Supplementary Methods.

### Differentially variable similar mean (DVSM) analysis

For the DVSM analysis, we included only genes that were expressed in at least half of the cells and had a large enough average expression level (> 5 normalized counts) in both conditions. For each gene, we defined the mean difference index between two gene *i,j* as 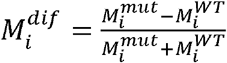 wher 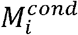 is the average expression of gene *i* in condition *cond*. Genes with mean difference index smaller than 0.05 (which corresponds to 10% difference) were considered as similar mean genes, and similar trends the results we present were also obtained with a cutoff of 0.1.

Based on the 3-fit parameters, each gene in a given condition was classified as either pure Gaussian or mixed. A gene was defined as pure Gaussian if the Gaussian fraction coefficient according to the fit was larger than 0.9. For similar mean genes we compared their variability between conditions according to their classification:

1. If both genes were classified as pure Gaussians, we used a bootstrap approach to calculate the confidence interval (CI) for the ratio between their variances. For the bootstrap calculation we sampled 10,000 instances for each condition. For each pair of instances, we calculated the variability ratio between the two conditions. If the 95% CI was larger (/smaller) than 1 the gene in the second (/first) condition was considered more variable.
2. If the gene is pure Gaussian in one condition and had a mixed profile in the second with Gaussian fraction smaller than 0.8, which corresponded to a distribution that contains very small (i.e. zeros) and/or very large extreme values, then it was considered more variable in the second condition.

#### CV calculation

We observed that the CV in our data is very sensitive to removal of 1-2 extreme data points for a gene (Figure S4E). This sensitivity may result in an effect of as high as 100% on the CV estimation. We therefore refined the CV measure to what we call trimmed-CV, such that the extreme top and bottom 5% of the data for every gene is eliminated from the analysis. This results in a more robust estimation.

#### DVSM based on CV

Similar to the comparison based on the modeling approach in the case of two pure Gaussians (type I), we used a bootstrap approach to calculate the confidence interval and the related p-values for the ratio between the trimmed-CV in mutant vs WT cells for every gene with similar mean. In addition, we used a permutation test mixing the cells identity between WT and mutant to calculate the corresponding p-value (Table S4).

### Variability rank score and significant differential-variability (DV)

In order to calculate the variability rank score we first computed for every gene the trimmed coefficient of variation (CV). We then sorted all genes by their mean expression level, and calculated for every group of 150 consecutive genes the relative rank of their trimmed-CV compared to the other genes in the group and normalized to values between 0 and 1. The number of genes for averaging was chosen based on stability of the resulting variability rank score (see Figure S4F). This resulted in every gene having 150 different rank scores, besides the extreme genes with the largest/smallest expression levels which were removed from further analysis. Next, for every gene we took the average of the 150 rank scores as the final variability rank score. For significance levels, we calculated the standard deviation of the variability rank difference of the two conditions, and used genes that had an absolute difference larger than 1 or 2 stds.

To define differentially-variable (DV) genes, we first calculated the standard deviation of the distribution of the difference of the variability rank scores between the two conditions. DV-genes were defined as genes that had a variability rank score difference of 1 or 2 standard deviations from the mean difference between the two conditions.

### Motif and gene annotation analysis

To search for enrichment of DNA sequence motifs as well as performing GO analysis we used HOMER (http://homer.ucsd.edu/homer/). We used the findMotifs.pl function for promoter regions from −500 to +100 relative to the TSS. To avoid bias towards more expressed genes, we used the background list option with the proper background list which contained only genes that were expressed above a defined threshold.

For the HTT interaction partners enrichment analysis we downloaded data from BioGRID (https://thebiogrid.org/).

### Image analysis

We used CellProfiler [65] to analyze the images that contained nuclei and aggregates. We built a separate pipeline to identify each. Identified aggregates were then linked to the closest nucleus in the image. For more details see Supplementary Methods.

## Declarations

### Availability of data and materials

The datasets generated and analyzed during the current study are available in the GEO repository (GSE138525). All custom code used in the current study is available from the authors upon a reasonable request.

### Authors’ contributions

M.S. and S.S. performed cell culture experiments and analyzed the data. W.O. performed siRNA screen and analyzed the data. M.N-R. performed cell culture experiments. X.X. and M.L.P. derived the 180Q isogenic cells, L.M.E. derived the 72Q isogenic cells, X.S., C.L., J.Q.P., N.S. and F.Z. derived the CHD8^+/-^ isogenic cells. C.C.H., X.A., S.K.S., and J.Z.L. performed the scRNA-Seq experiments. M.S. and E.M. designed the experiments and wrote the manuscript.

## Acknowledgments

Alzheimer-related data were provided by the Rush Alzheimer’s Disease Center, Rush University Medical Center, Chicago. Data collection was supported through funding by NIA grants P30AG10161, R01AG15819, R01AG17917, R01AG30146, R01AG36836, U01AG32984, U01AG46152, the Illinois Department of Public Health, and the Translational Genomics Research Institute. The work was supported in part by grants P30AG10161, R01AG15819, R01AG17917, and U01AG61356. ROSMAP data can be requested at www.radc.rush.edu.

