## Supplemental Figures and legends for "Pluripotent stem cell derived models of neurological diseases reveal early transcriptional heterogeneity"

### Supplementary Figures and Figure legends

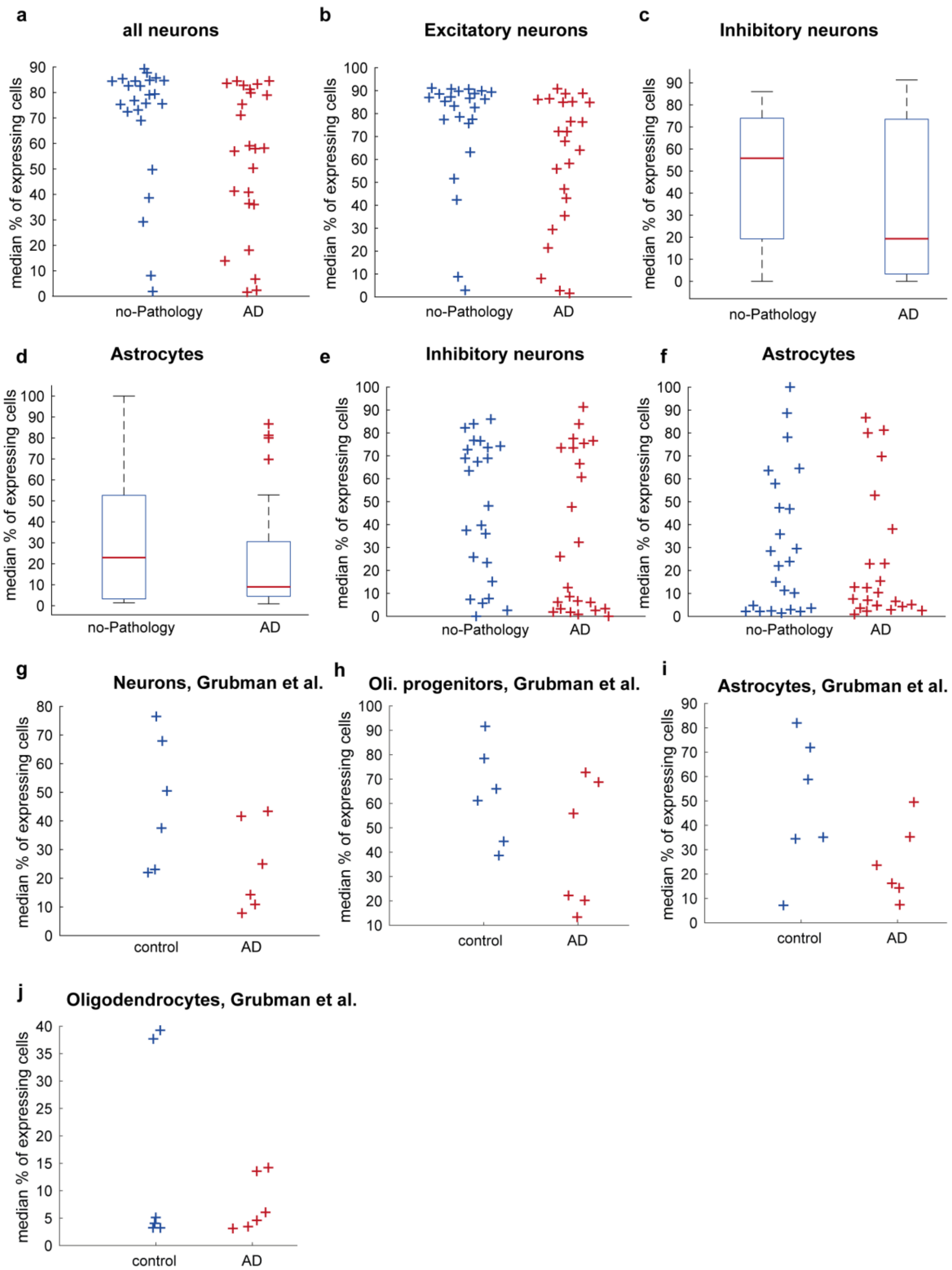


**Figure S1. Transcriptional heterogeneity in AD adult neurons. Related to Figure 1.** (A-F) The median percentage of expressing cells of the most 200 expressed genes for all adult neurons (A), excitatory neurons (B), inhibitory neurons (D) and astrocytes (F) in healthy and AD subjects from Mathys et al. and the corresponding boxplots for inhibitory neurons and astrocytes (C and E, respectively). (G-J) The median percentage of expressing cells of the most 100 expressed genes for all adult neurons (G), oligodendrocyte progenitor cells (OPC, H), astrocytes (I) and oligodendrocytes (J) in control and AD subjects from Grubman et al.

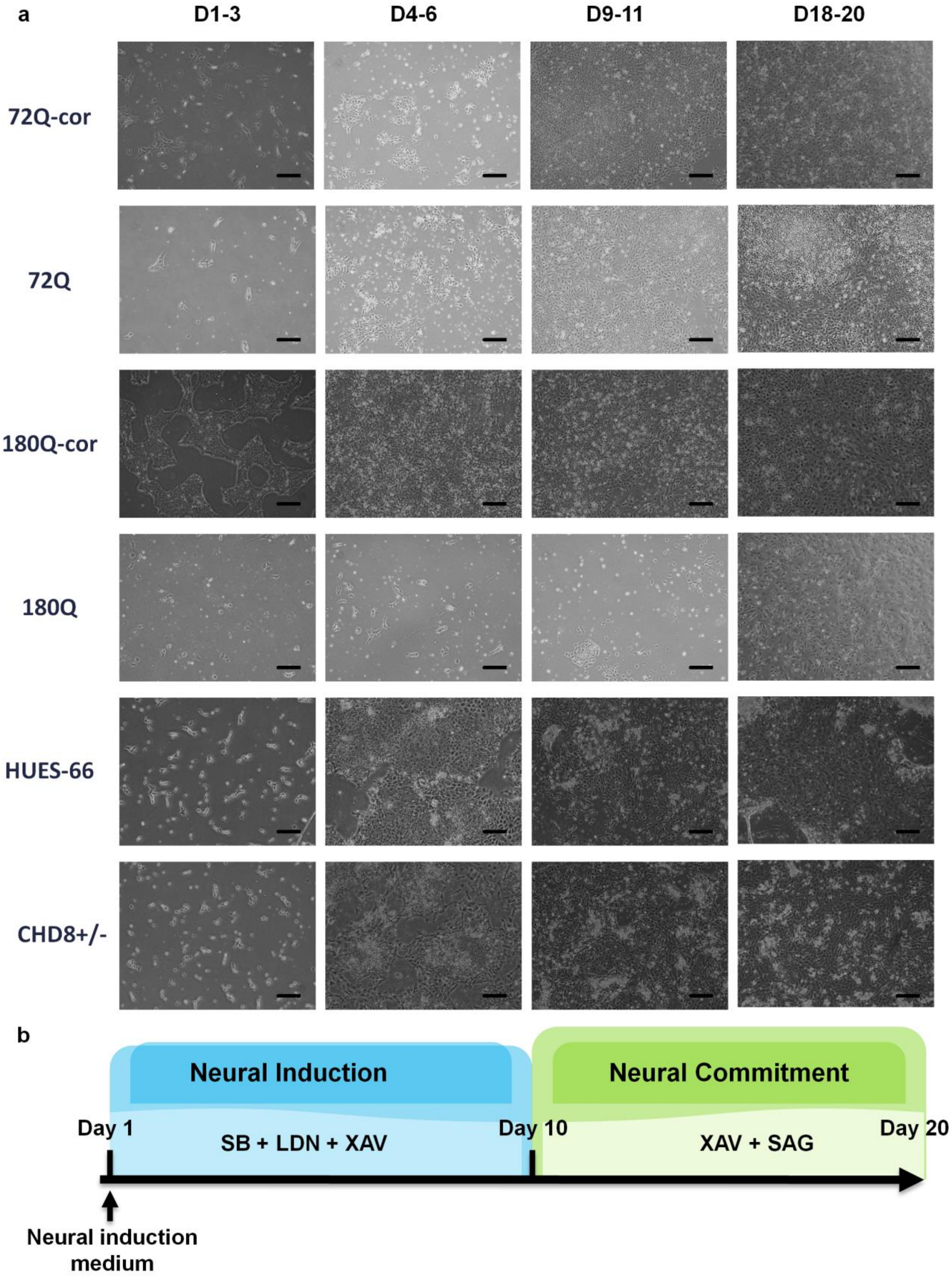
a
D1-3
D4-6
D9-11
D18-20
72Q-cor
72Q
180Q-cor
180Q
HUES-66
CHD8+/-
b
Neural Induction
Neural Commitment
Day 1
SB + LDN + XAV
Day 10
XAV + SAG
Day 20
Neural induction medium

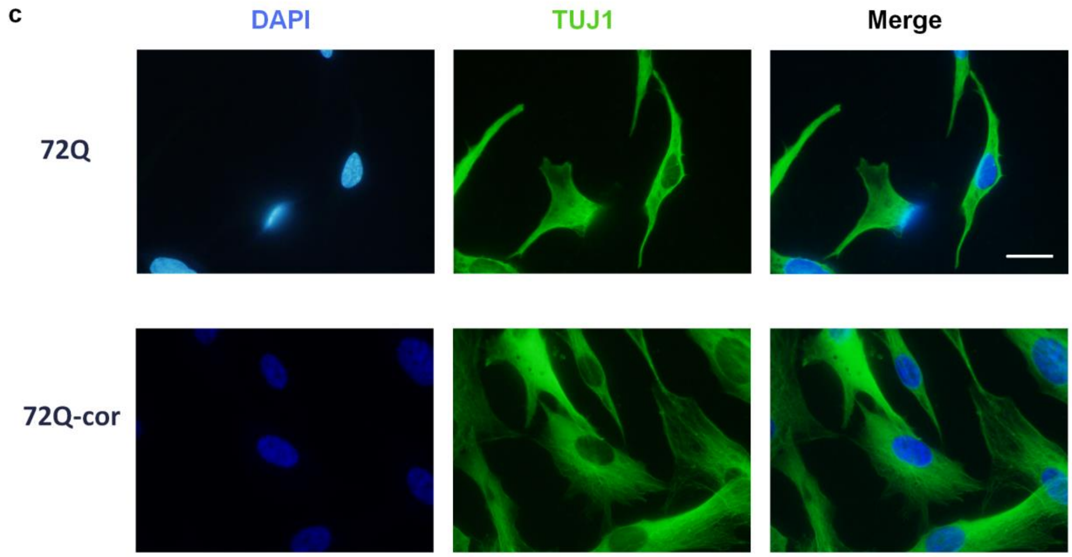


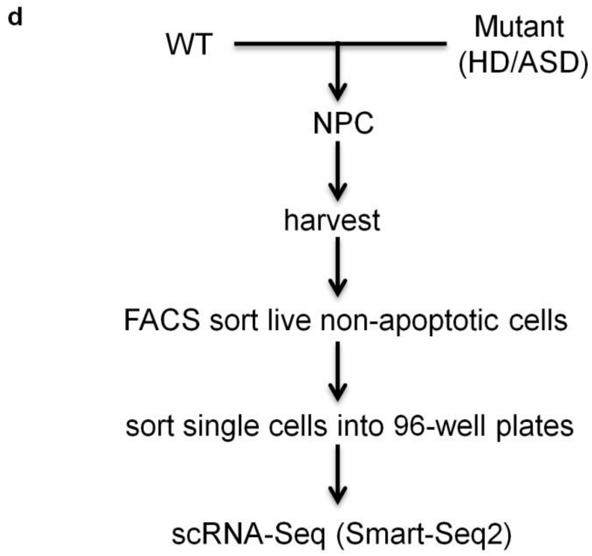


**Figure S2. Experimental setup of scRNA-Seq of neuronal progenitor cells in culture. Related to Methods.** (A) During differentiation, cells gradually acquire neuronal identity. Shown are days selected stages of the protocol. Scale bars = 50 µm. (B) Overview of the neuronal differentiation protocol of pluripotent stem cells. (C) NPCs express the neuronal marker TUJ1. Shown are example of the mutant and corrected 72Q cells. Scale bars = 10 µm (D) Experimental pipeline.

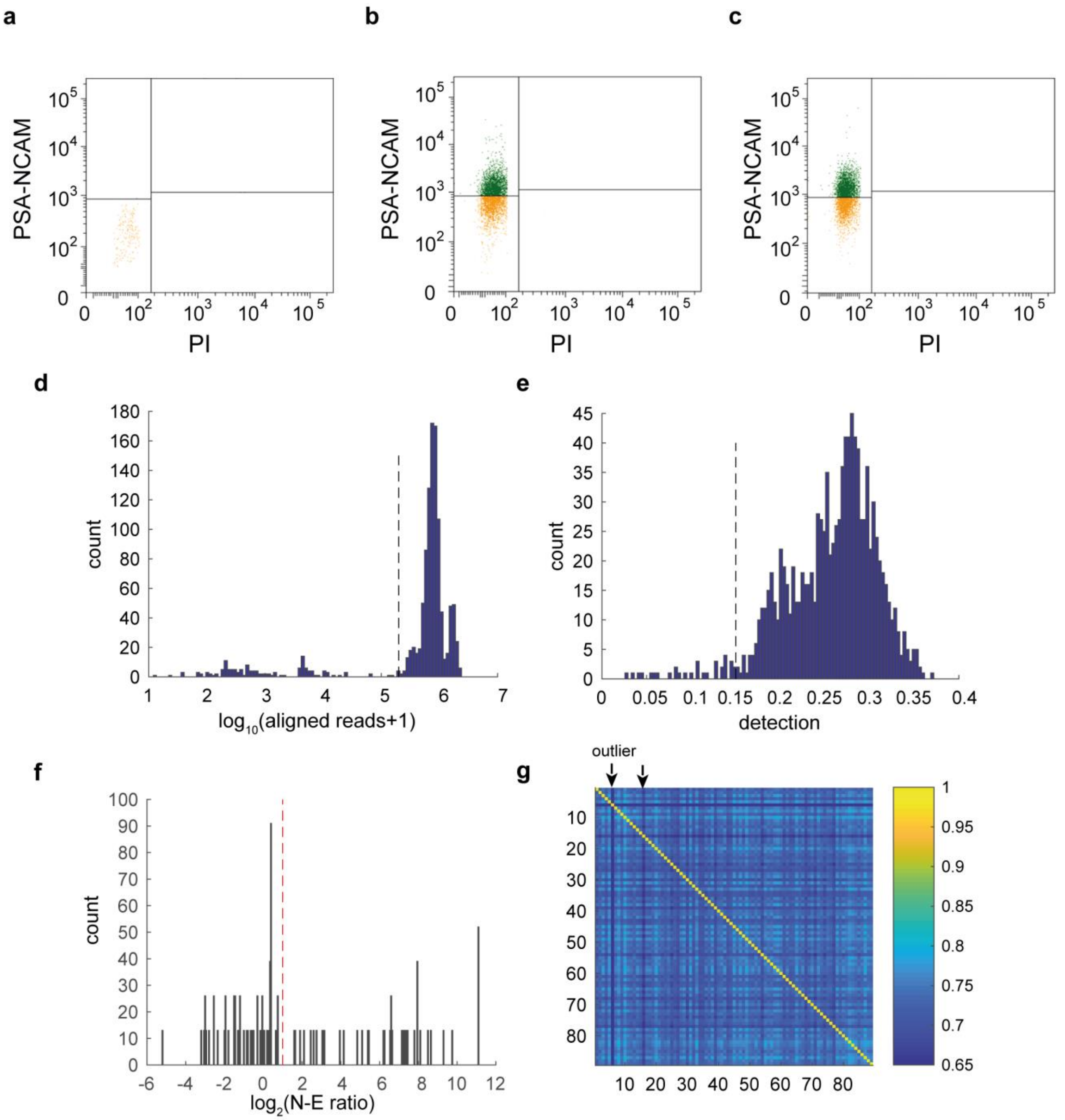


**Figure S3. Experimental and computational selection of high-quality NPC for analysis. Related to Methods.** (A-C) Cells are FACS-sorted using Annexin-V and PI to filter out apoptotic and dead cells, respectively, and to select for NPCs based on the expression of PSA-NCAM. Dark green represents cells which are positive for PSA-NCAM and negative for both Annexin-V (not shown) and PI. (A) hESC serve as negative controls for PSA-NCAM. (B-C) A large percentage of the mutant (here 72Q) and WT (72Q-corrected) NPCs, respectively, are intact and enriched for PSA-NCAM. (D) Histogram of total number of reads aligned to the transcriptome in all cells. (E) Histogram of percentage of detected genes in every cell. (F) Histogram of neural-to-embryonic **n-e** ratio of the cells (see Methods for definition). In (D-F) black dashed line represents lower threshold for filtering out cells. See Methods for more details. (G) Example of Spearman correlation between every 2 cells in the 72Q iPSC-derived NPCs. Arrows point at outliers.

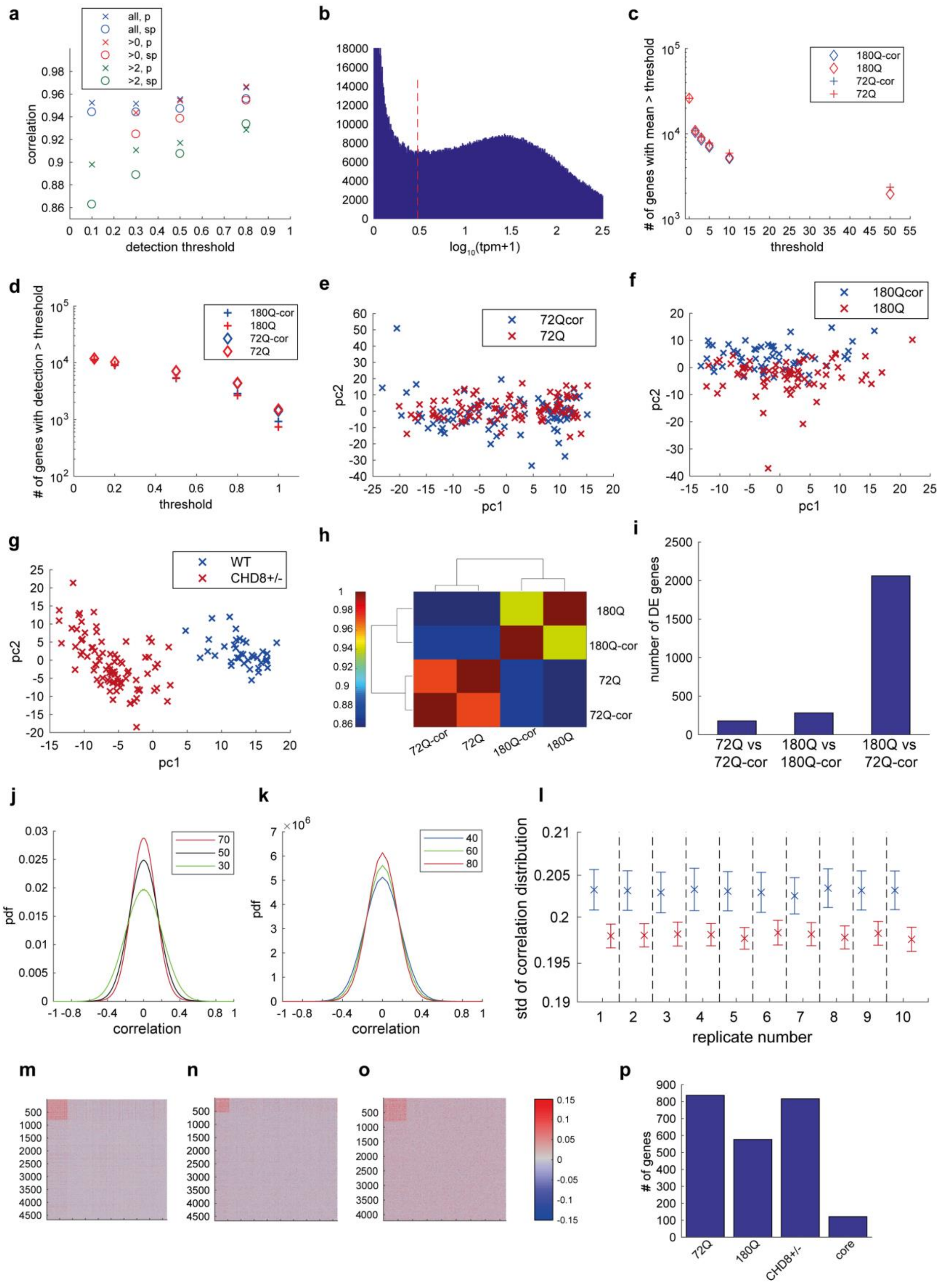

**Figure S4. Correlation between single-cell and bulk data and HD transcriptional changes based on average expression level and pairwise gene correlations. Related to Figure 2.** (A) Correlation of bulk data and average expression of single-cell data when including all values (“all”), only positive value (“>0”) and only reliable values (“>2”) using threshold from (B). "sp" stands for Spearman correlation and "p" stands for Pearson correlation. (B) Distribution of normalized expression levels in single-cell data. Red dashed line marks threshold used for detection. (C) Number of genes detected in single-cell data above different expression thresholds. (D) Number of genes detected in single-cell data above different thresholds of fraction of cells. (E-G) PCA of the 72Q, 180Q and $CHD8^{+/-}$ isogenic system, respectively. (H) Spearman correlation between mean expression in HD and WT cell lines. (I) Number of differentially expressed (DE-) genes in the HD isogenic systems and between the non-isogenic pair of 180Q mutant and the 72Q-corrected. (J) The correlation distribution depends on the number of cells; The larger the number of initial pool of cells, the smaller the correlations are in absolute value (see Methods). Shown are the distribution of pairwise gene correlations as a function of the number of cells used for the calculation out of the initial pool. The results shown are based on the average of 100 random samplings from the initial population of cells. (K) Given the same size of initial pool of cells, the correlation distribution depends on the number of cells drawn for the analysis; The larger the number of drawn cells, the smaller the correlations are in absolute value (see Methods). The results are based on summation of 50 random samples for each number of initial pool of cells. Results in (J) and (K) are shown for the 72Q-cor cell NPCs. (L) The stds of the correlation distributions in 10 replicates of random initial pools averaged over 1000 random selections of 30 cells in the 72Q isogenic system. Blue, WT; red, mutant. (M-O) Cluster analysis based on the difference of all pairwise correlations between WT and mutant cells in 72Q, 180Q and $CHD8^{+/-}$ (M-O, respectively) isogenic systems. (P) Number of genes in the major cluster (sub-network) of genes with a decreased correlation in the mutant cells.

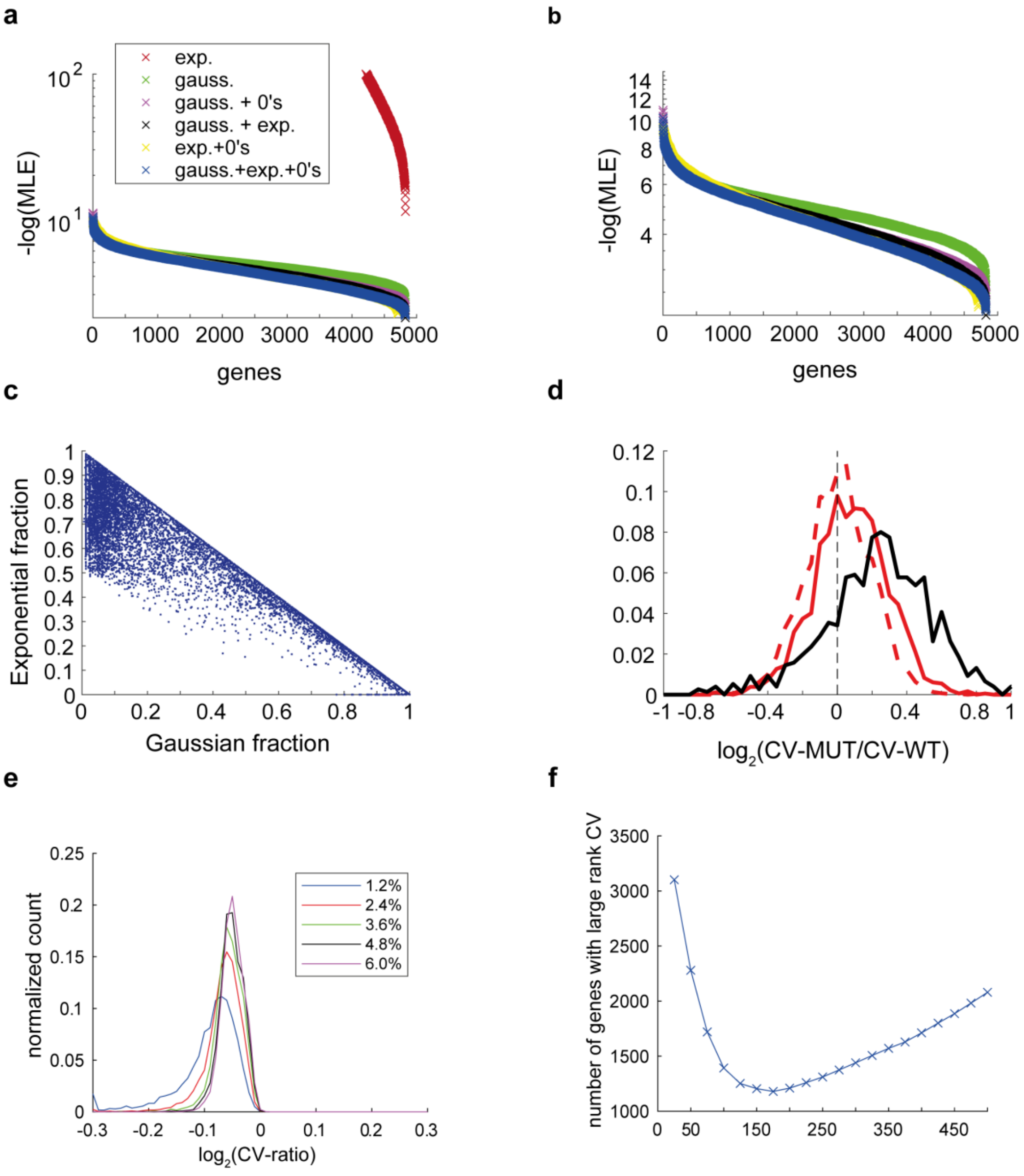


**g** **NKX2-1**

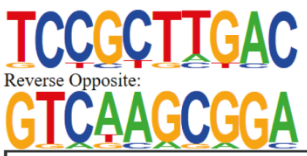


| | |
|---|---|
| p-value: | 1e-10 |
| log p-value: | -2.429e+01 |
| Information Content per bp: | 1.844 |
| Number of Target Sequences with motif | 13.0 |
| Percentage of Target Sequences with motif | 11.30% |
| Number of Background Sequences with motif | 29.0 |
| Percentage of Background Sequences with motif | 0.72% |

**h** **GMEB1**

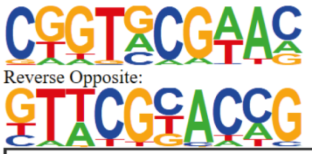


| | |
|---|---|
| p-value: | 1e-10 |
| log p-value: | -2.444e+01 |
| Information Content per bp: | 1.700 |
| Number of Target Sequences with motif | 18.0 |
| Percentage of Target Sequences with motif | 18.18% |
| Number of Background Sequences with motif | 89.0 |
| Percentage of Background Sequences with motif | 2.18% |

**Figure S5. Model fit for gene expression distribution and characterization of differentially-variable genes. Related to Figures 3 and 4.** (A-B) Shown is the (sorted) average log-likelihood of the best fit for 6 different models (see Methods for more details) in the 180Q-corrected cell line: exponential distribution, Gaussian distribution, mixture of Gaussian and uniform zeros distributions, mixture of Gaussian and exponential distributions, mixture of exponential and uniform zeros distributions and mixture of Gaussian, exponential and uniform zeros distributions. The mixture model which uses 3 distributions outperforms the rest of the models. (B) As in (A), plotted with better resolution for lower values. (C) The Exponential and Gaussian fractions of the best fir of the 3 distributions mixture model of all expressed genes from the 6 cell lines used in the this study. (D) Same as Figure 3I after removal of genes in suspicious chromosomal regions. The distribution of the log ratio of the CV between mutant and WT for the three isogenic systems. p-value for the $CHD^{+/-}$ system in this analysis is << $10^{-60}$ (Wilcoxon signed rank test). 72Q: dashed red line, 180Q: red solid line, $CHD8^{+/-}$: black solid line. (E) The CV is more stable after trimming the outliers. Shown is the distribution of the log2 ratio of the CV before vs after exclusion of additional 1.2% of the extreme low and high expression values. Extreme 1.2% - blue, 2.4%, 3.6%, 4.8%, 6% in red, green, black and purple, respectively. (F) Rank score stability as a function of the size of the running window. Shown is the number of genes with larger noise (measured by CV) in their calculated rank score (see Methods for more details) above noise thresholds of 0.1. The optimum window size, where the number of genes that succeed the noise threshold is minimal, is achieved around 150 points. (E-F) The enriched known binding motifs in the promoter regions of DV-genes. (G) DV-genes, which are more variable in 180Q mutant cells, are enriched for NKX2-1 binding motif. (H) DV-genes, which are more variable in 180Q-corrected cells, are enriched for GMEB1 binding motif.

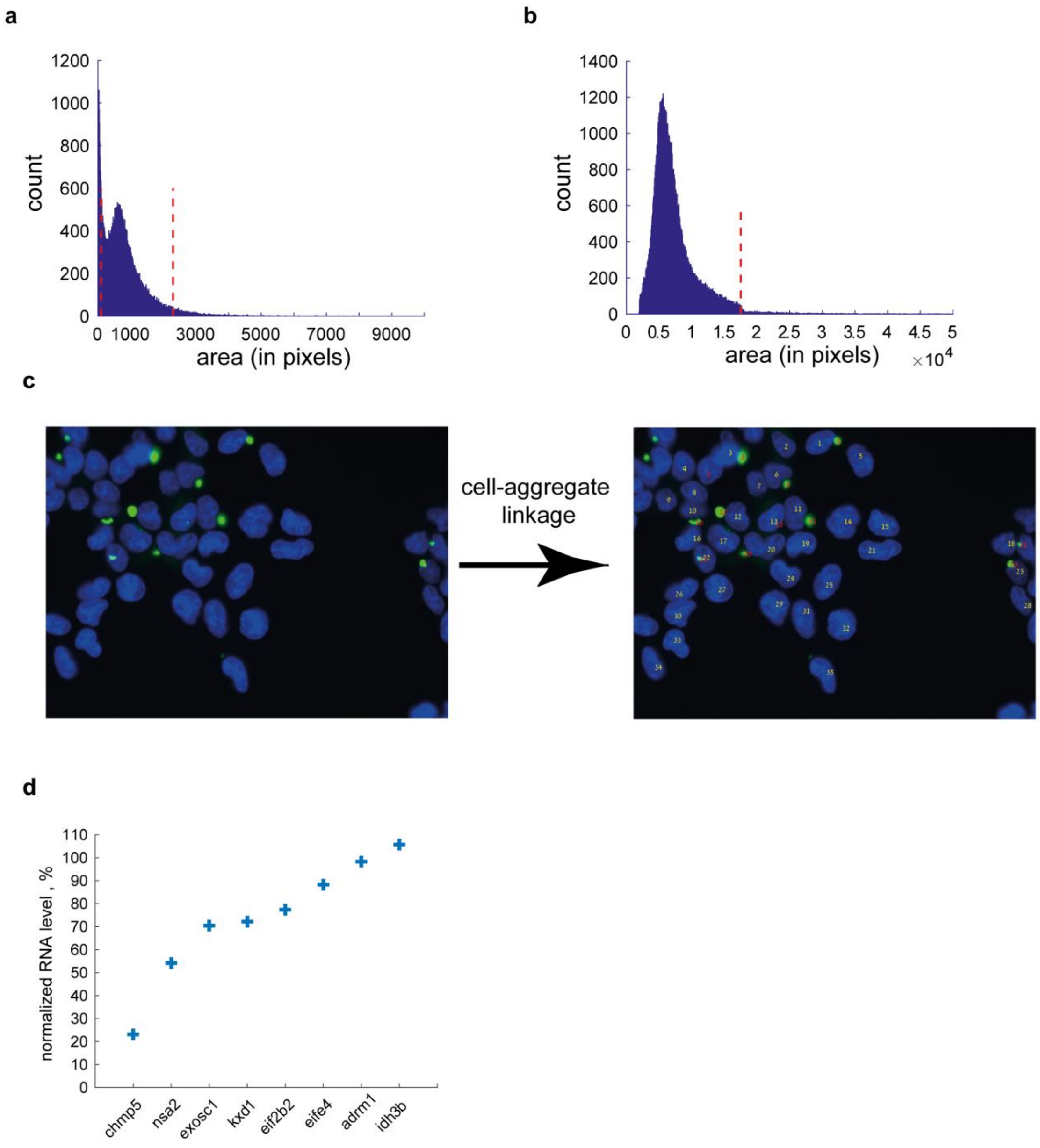


**Figure S6. Image analysis of aggregate functional assay. Related to Figure 5.** (A-B) Filtering out suspicious results. **a** Distribution of nuclei area across all images. Nuclei are filtered out if they are too large or too small (thresholds marked by the red dashed line). (B) Distribution of aggregates size across all images. Aggregates are filtered out if they are too large (threshold marked by the red dashed line). (C) At the last stage, identified nuclei and aggregates are linked to each other based on their distance. Numbers on nuclei represent the cell numbering and numbers on aggregates represent the cell number to which the aggregates are assigned. (D) The percentage of RNA levels in different gene knockdown in NPCs. Expression levels are normalized to ACTβ and to the median expression levels in non-treated (WT) and scrambled samples.
